# Image retrieval based on closed-loop visual–semantic neural decoding

**DOI:** 10.1101/2024.08.05.606113

**Authors:** Ryohei Fukuma, Takufumi Yanagisawa, Hidenori Sugano, Kentaro Tamura, Satoru Oshino, Naoki Tani, Yasushi Iimura, Hui Ming Khoo, Hiroharu Suzuki, Huixiang Yang, Takamitsu Iwata, Madoka Nakajima, Shinji Nishimoto, Yukiyasu Kamitani, Haruhiko Kishima

## Abstract

Neural decoding via the latent space of deep neural network models can infer perceived and imagined images from neural activities, even when the image is novel for the subject and decoder. Brain-computer interfaces (BCIs) using the latent space enable a subject to retrieve intended image from a large dataset on the basis of their neural activities but have not yet been realized. Here, we used neural decoding in a closed-loop condition to retrieve images of the instructed categories from 2.3 million images on the basis of the latent vector inferred from electrocorticographic signals of visual cortices. Using a latent space of contrastive language-image pretraining (CLIP) model, two subjects retrieved images with significant accuracy exceeding 80% for two instructions. In contrast, the image retrieval failed using the latent space of another model, AlexNet. In another task to imagine an image while viewing a different image, the imagery made the inferred latent vector significantly closer to the vector of the imagined category in the CLIP latent space but significantly further away in the AlexNet latent space, although the same electrocorticographic signals from nine subjects were decoded. Humans can retrieve the intended information via a closed-loop BCI with an appropriate latent space.

## Introduction

Imagery-based information retrieval via brain–computer interfaces (BCIs) will extend information retrieval and communication capabilities for humans, including patients with various neurological diseases. Although some invasive BCIs have demonstrated clinical feasibility in supporting the communication of paralyzed patients, most invasive BCIs rely on motor cortical activities to control external devices^1-4^. These BCIs have limited applicability to patients with deteriorated motor cortical activity, such as those with severe amyotrophic lateral sclerosis (ALS)^5^. BCIs that output the intended information independent of motor cortical activity could lead to BCI-based communication aids for those patients^6^. In addition, such BCI-based information retrieval could improve the quality of semantic search, which is often difficult due to the ambiguity and complexity of queries^7,8^. The ability of imagery-based BCIs to retrieve information may enhance information retrieval and communication in humans but has not yet been realized.

Visual–semantic information encoded in visual cortices is beneficial for BCIs. Previous studies have shown that various visual and semantic attributes of the images perceived by a subject can be inferred (decoded) from the subject’s visual cortical activity recorded by functional magnetic resonance imaging (fMRI) scans and electrocorticography (ECoG) signals^9-13^. In particular, using deep neural network (DNN) models, which encode visual and semantic attributes of various images in their latent vector, neural decoders trained with these latent spaces can decode neural activities to infer images that have not been used in the training of the decoder^9,10,12,14^. In other words, zero-shot learning to infer visual–semantic attributes is possible with an appropriate latent space. In addition, such a neural decoder trained to infer the perceived images is also capable of inferring some images that the same subjects imagined^9,15^, suggesting that the trained decoder learnt the common neural representation between imagery and perception^16^ and that imagery-based retrieval of the intended image is possible. However, low decoding accuracy makes it difficult to use for practical information retrieval and communication^17^.

Closed-loop control of the decoded images may allow accurate retrieval of the intended image by modulating the decoded images cognitively. Previous studies have demonstrated the closed-loop control of images decoded from neural activities^17-19^. Subjects could control two different superimposed images whose visibility ratios changed depending on the activity of the hippocampal neurons^19^ or fMRI scans of the visual cortex^18^ by imagining and attending one of the overlayed images. Attention and imagery could be the top-down mechanism to control such feedback images. Attention has been shown to alter visual cortical activities^20^ so that the decoded image becomes closer to the attended image^21^. In addition, it has been demonstrated that subjects can control feedback images to represent the intended meaning via imagery when the feedback images are determined on the basis of semantic vectors decoded from ECoG signals^17^. However, in this study, the semantic vector for each image was created from human annotations; the size of the image pool to be searched was therefore limited to a few thousand images, which was insufficient to include a variety of novel images. Taken together, these results suggest that, in a closed-loop condition, decoded images can be controlled by imagery and attention via visual neural decoding, although the retrieval of images novel for the subjects and decoder has not yet been realized. The combination of neural decoding, the latent space of the DNN model, and a large image pool enables the subject to retrieve various novel images via zero-shot learning. Here, it is hypothesized that, when an appropriate latent space is used, subjects can control a high-dimensional decoded latent vector encoding visual–semantic information through a closed-loop BCI to retrieve images of the intended meaning from a large image pool.

To demonstrate proof-of-concept information retrieval via BCIs, we developed a BCI that retrieves images from a dataset consisting of 2.3 million images on the basis of visual– semantic vectors decoded from human ECoG signals. To train the decoder, 1,200 images of natural scenes were presented to the electrode-implanted subjects (image perception task) to record ECoG signals from the occipital, temporal and parietal cortices, while the visual– semantic vector for each image was obtained by encoding the images with a DNN model. With image retrieval based on a visual–semantic vector decoded in real time, the subjects were presented with the retrieved (feedback) image (closed-loop condition) to control the feedback image via imagery so that the feedback image represented the instructed meaning (online task). With this task, the accuracy of following the instructions was compared between two latent spaces: those of contrastive language-image pretraining (CLIP)^22^ and AlexNet^23^ models. The CLIP model consists of an image encoder and a text encoder; both encoders are trained simultaneously by inputting the paired image and corresponding annotation to discriminate between correct and incorrect pairs^22^. The AlexNet model, whose architecture is a convolutional neural network, was trained to classify images into 1,000 categories^24^. This study utilized the latent space commonly shared by the image and text encoders of the CLIP model and the latent space spanned by unit responses in the middle layer (fc6) of the AlexNet model (AlexNet-fc6). In addition to the online task, another task (modulation task) was performed to evaluate how imagery affects the decoded vector, especially depending on the latent space; in this modulation task, the change in the decoded vector towards the vector of the imagined meaning was evaluated when the subject imagined an image while viewing another image representing a different meaning from the imagery.

## Results

### Neural decoding of the perceived images

First, five subjects who later participated in the online task performed an image perception task to create a decoder to infer latent vectors representing visual–semantic information of images from various semantic categories. These subjects were implanted with subdural electrodes on the occipital, temporal, and parietal cortices (Supplementary Fig. 1a and b, and Supplementary Table 1). In the image perception task, 1,200 images from 150 categories and 50 images from the other 50 categories were presented for 500 ms each to the subjects as training and test images for the decoder, respectively (Fig. 1a; also see Methods, Supplementary Fig. 1c and d). The stimuli images were encoded by either the CLIP image encoder model or the AlexNet model to acquire 512-dimensional vectors in the CLIP latent space (CLIP vector) or 1,000-dimensional vectors from units of the fc6 layers (AlexNet-fc6 vector) (see Methods). A ridge regression model (decoder) was trained to infer the latent vectors of the training images (1,200 images) from the standardized high-γ powers (high-γ features; 80–150 Hz; see Methods) in the 500-ms ECoG signals during the presentation of the corresponding stimuli images.

**Fig. 1.**
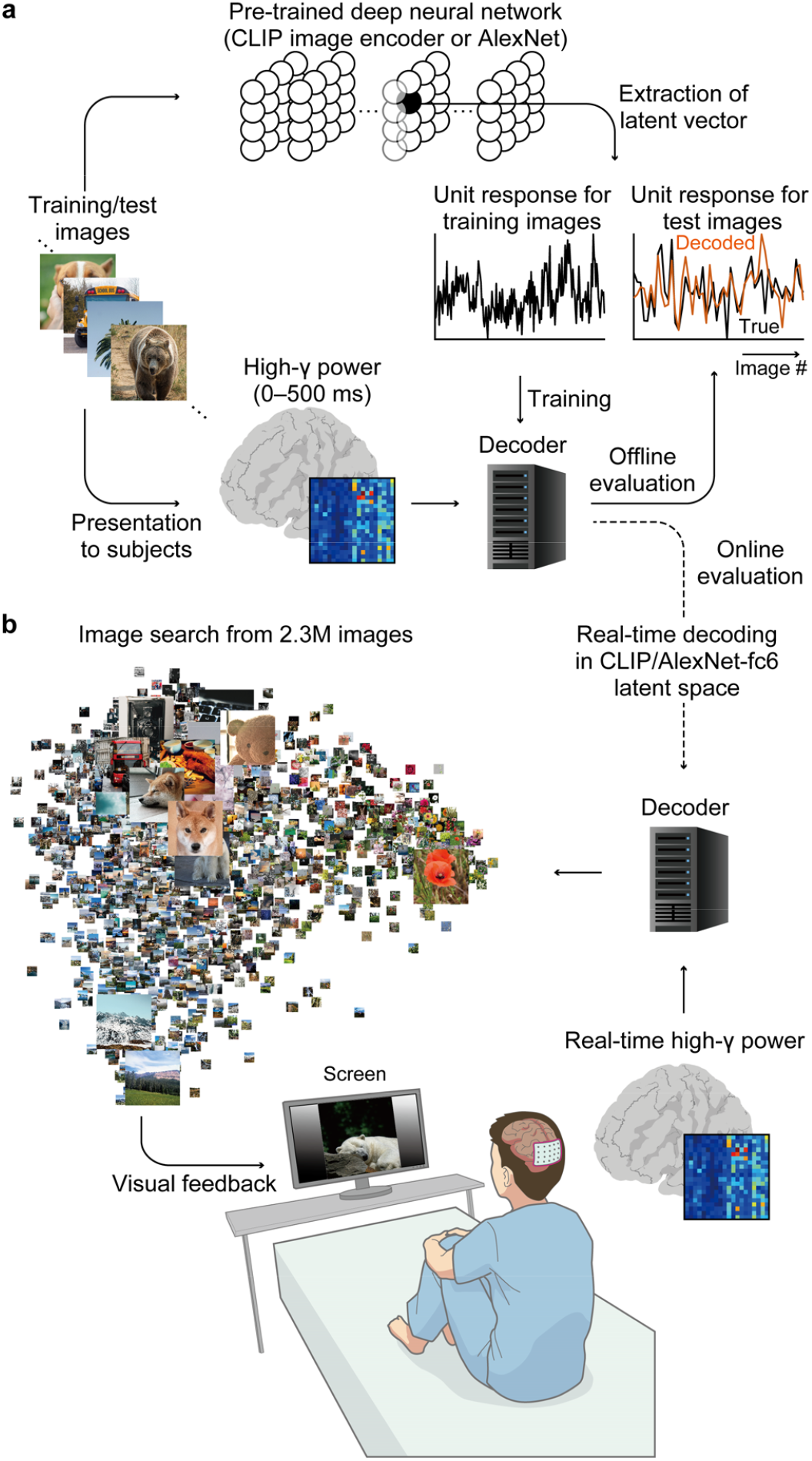
Decoder training and outline of the online task. (a) Under the recording of ECoG signals, each subject participated in one or more training and test sessions of the image perception task, in which 1,200 GOD training images and 50 test images were presented to the subjects. The images were fed into a CLIP image encoder model or AlexNet model to acquire latent vectors (CLIP vectors or AlexNet-fc6 vectors). A linear decoder model was trained so that these latent vectors of the training images could be decoded from the high-γ features (standardized high-γ powers) during the presentation of the corresponding training images (0–500 ms). To evaluate the decoding accuracy in the offline condition, the decoder was applied to the high-γ features of the test images. (b) Using the decoder acquired in (a), five subjects participated in the online task. From their real-time high-γ features, a latent vector was inferred in the CLIP or AlexNet-fc6 latent spaces every 250 ms; a feedback image was searched from the Unsplash image dataset or the GOD training image dataset on the basis of the highest cosine similarity (or Pearson’s correlation coefficient for EB01 and EB02) with the inferred vector.

The performance of the trained decoder was evaluated by decoding the high-γ features while presenting the 50 test images. The decoding accuracies were measured by the accuracy of identifying the corresponding image via the inferred latent vector from other test images (scene-identification accuracies) (see Methods). For all the subjects, the scene-identification accuracies were significantly higher than the chance level estimated by permutation (*p* < 0.05, one-sided paired *t* test) for both the CLIP latent space (72.50 ± 9.62% (mean ± 95% confidence intervals (CIs), *n* = 5); Table 1) and the AlexNet-fc6 latent space (74.63 ± 7.88%). Notably, the accuracy was assessed with images whose categories are novel to the decoder because the 50 categories in the test images did not overlap with the 150 categories in the training images. Hence, the latent vectors corresponding to the various novel images were successfully decoded in a zero-shot manner from the high-γ features of the temporal, parietal and occipital cortices.

**Table 1.**
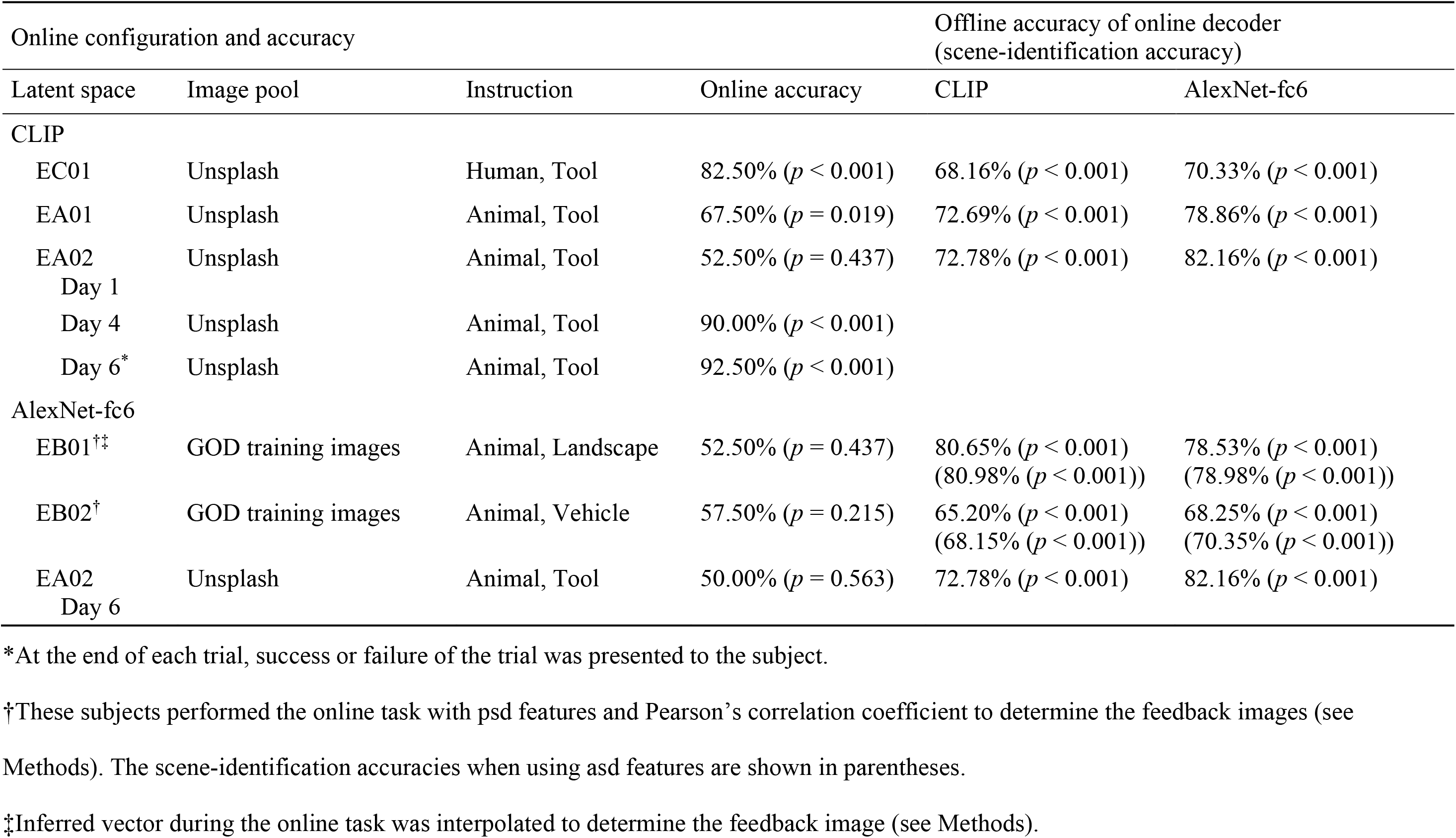
Summary of individual online/offline accuracy.

### Image retrieval in the CLIP latent space

Using the trained decoder with the CLIP vector, three subjects participated in the online task (Table 1). In the task, the subjects were instructed to control the feedback image shown on a computer screen via visual imagery so that the feedback image represented the instructed meaning (closed-loop condition; Fig. 1b and Supplementary Fig. 1e). Each trial started with one of two instructions, followed by 2.5 s of a blank screen and 32 feedback images with intervals of 250 ms. The set of instructions was determined prior to the task on the basis of each subject’s preference by asking the subject after several minutes of free-running feedback, during which the subjects freely controlled the feedback images (see Methods). Using the decoder trained from the high-γ features in the image perception task, each feedback image was selected from an image pool based on the highest cosine similarity between the latent vectors of the pooled images and the inferred vector in real time.

The subject EC01, whose electrodes were implanted over the parietal, temporal, and occipital lobes (Fig. 2a), was instructed to show an image of “tool” or “human” from an image pool that had approximately 2.3 million images of various contents (Unsplash^25^). The images used to train the decoders were not included in the image pool (see Methods). Fig. 2b shows feedback images presented to the subject during the first trial, whose instruction was “tool” (for the second trial with “human” instruction, see Supplementary Fig. 2a). The feedback images of the trials predominantly showed “tool” content. To evaluate how much the inferred CLIP vector became closer to the instructed meaning, the cosine similarity of the inferred CLIP vector to the instruction vectors for “tool” or “human”, which were acquired by the CLIP text encoder model, was assessed. The cosine similarity to the target instruction vector (“tool”, red line in Fig. 2b) was greater than that to the nontarget instruction vector (“human”, black line) in most of the 32 feedback images. This trial was considered successful because the average cosine similarity among the 32 feedback images was greater for “tool” than for “human”. On average over all 40 trials, the similarity to the target instruction vector was greater than that to the nontarget instruction vector, especially in the latter half of the trials (Fig. 2c left and Supplementary Fig. 2b). Consequently, EC01 achieved 82.50% online accuracy (Fig. 2c right and Table 1), which was significantly greater than the chance level (*p* < 0.001, one-sided binominal test, *n* = 40).

**Fig. 2.**
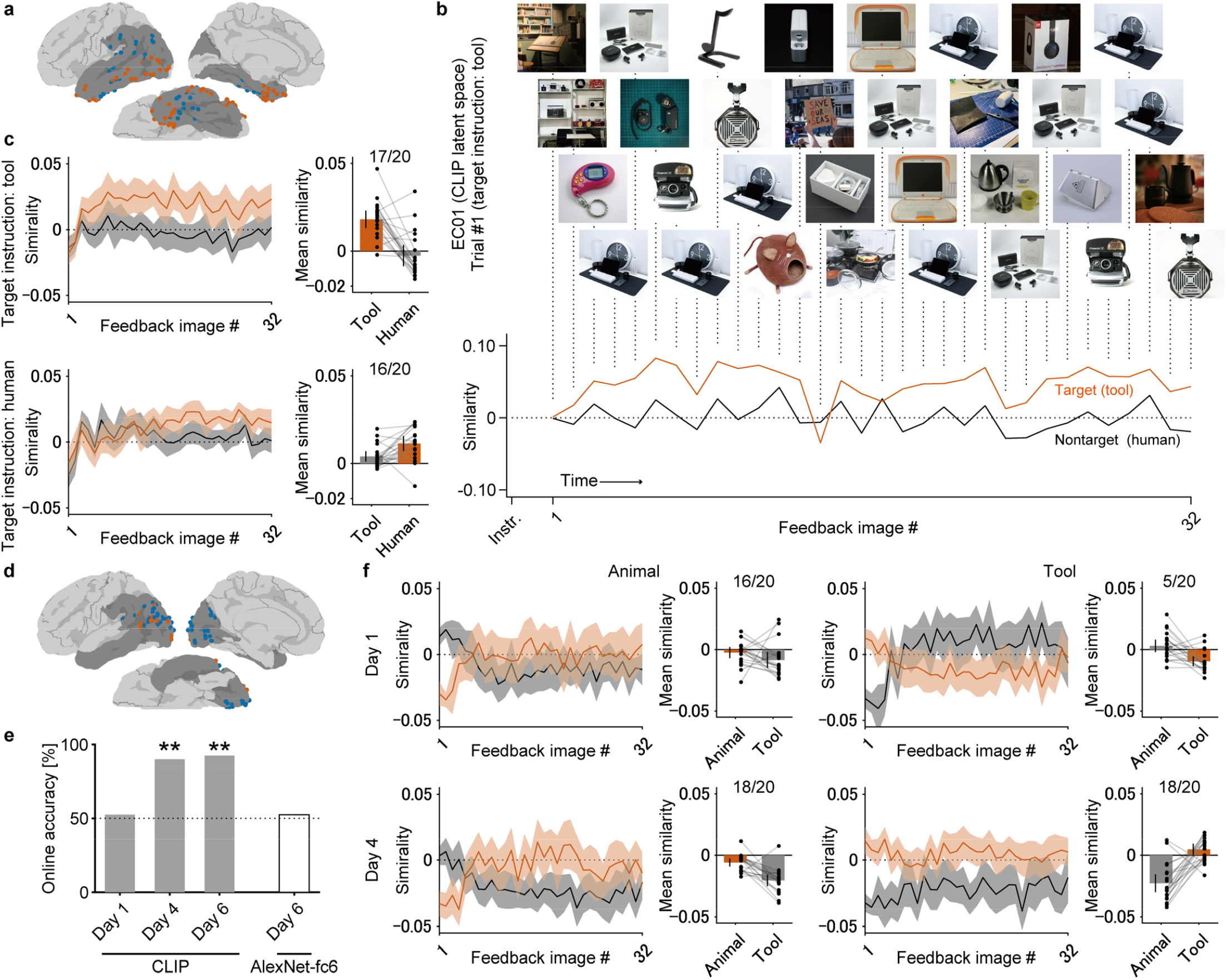
Closed-loop online task using the CLIP latent space by EC01 and EA02. (a) The subdural electrode location of EC01 is shown on the normalized brain. The electrodes on the left and right hemispheres are shown in red and blue, respectively. The cortical area marked with a darker colour denotes the regions where the subdural electrodes were located. (b) A representative trial (trial #1) of EC01 is shown with feedback images (top) and the cosine similarity of the inferred vector to the target/nontarget instruction vector. (c) Line graph (left) showing the trial-averaged cosine similarity between the inferred vector and target/nontarget instruction vector, with the coloured area denoting the 95% CI, where the bars (right) show the trial– and feedback-image-averaged cosine similarities with the corresponding 95% CIs. Individual values are represented with dots. On the basis of these average similarities in the bar graph, the success/failure of the trial was determined; the number of successes is shown at the top of the graph. (d) The subdural electrode location of EA02 is shown in the same format as (a). (e) Online accuracies of the closed-loop online task for EA02 are shown with bars. For the task with the CLIP latent space on Day 6, the success/failure of each trial was displayed at the end of the trial. **Bonferroni-adjusted *p* < 0.01, one-sided binominal test. (f) Similarity to the target/nontarget instruction vectors for EA02 on Days 1 and 4 is shown in the same format as (c).

In addition to EC01, two other subjects (EA01 and EA02) participated in the same online task. EA01 achieved a significant online accuracy of 67.50% (Table 1, *p* = 0.019, *n* = 40) for two instructions (“animal” and “tool”). On the other hand, EA02 participated in the task multiple times (sessions) with “animal” and “tool” instructions. On Day 1, the accuracy was 52.50% (Bonferroni-adjusted *p* = 1.312, *n* = 40); however, on Day 4, the accuracy reached 90.00% (Bonferroni-adjusted *p* < 0.001, *n* = 40) (Fig. 2d and e). The similarity to the target/nontarget instruction vector (Fig. 2f) showed that EA02 failed to increase the similarity to the target instruction during tool trials on Day 1 but succeeded on Day 4. The increase in the accuracy suggests that training to control the feedback images in the closed-loop condition improves accuracy. Finally, on Day 6, the inferred instruction (“animal” or “tool”) was presented on the screen at the end of each trial, with success/failure colour-coded so that the subject’s intention for each trial was clearly visible; with this setting, the accuracy reached 92.50% (Bonferroni-adjusted *p* < 0.001, *n* = 40; Supplementary Fig. 3), demonstrating a communication of intent of 0.6157 bits in 8 s of each feedback trial (0.0770 bps). These results demonstrate that BCIs using the CLIP vector inferred from ECoG signals enable subjects to retrieve images of their intentions from large image pools and to output their intentions in closed-loop conditions.

**Fig. 3.**
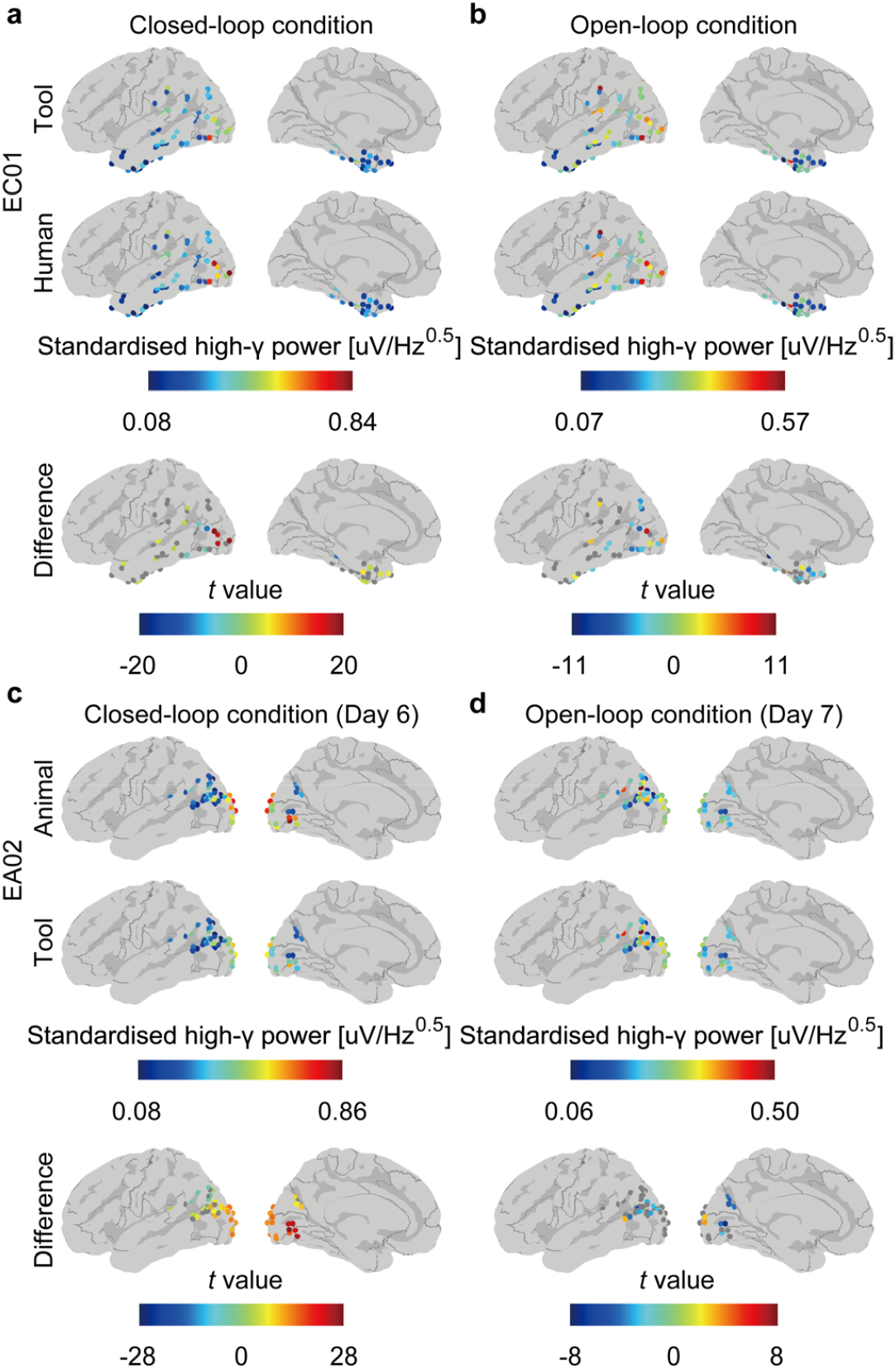
Cortical activities during the online task. (a, b) Trial and feedback image averages of high-γ features (standardized high-γ powers) used to determine the feedback images during the online task of EC01 with (a) the closed-loop condition and (b) the open-loop (eyes-closed) condition are visualized on a normalized brain surface with colour-coding. Differences in the high-γ features between the tool instructions and the human instructions were evaluated with two-sided Welch’s *t* test to visualize significant differences at each electrode (FDR-adjusted *p* < 0.05). (c, d) High-γ features for EA02 were also visualized in the same format as in (a) and (b). For the closed-loop condition, the activity on the nearest day (Day 6) as that of the open-loop condition (Day 7) was visualized.

In addition to the online accuracy that was evaluated on the basis of the given two instructions, the accuracy of identifying the instruction of each trial from the other six instruction candidates (retrieval accuracy) was evaluated in a two-choice manner (see Methods). The retrieval accuracy was 81.25% for EC01 (*p* < 0.001, *n* = 40, one-sided paired *t* test against chance level estimated by permutation) and 45.42% for EA01 (*p* = 0.815, *n* = 40); those of EA02 on Days 1, 4, and 6 were 25.42% (Bonferroni-adjusted *p* = 3.000, *n* = 40), 61.25% (Bonferroni-adjusted *p* = 0.014, *n* = 40), and 69.17% (Bonferroni-adjusted *p* < 0.001, *n* = 40), respectively.

### Cortical activities during the online task

The high-γ features (standardized high-γ powers) of the ECoG signals used to determine the feedback images were compared between two instructions to assess which cortical region contributed to closed-loop control. For EC01, the high-γ feature showed high values in the visual area during the online task, with a significant difference between the two instructions (FDR-adjusted *p* < 0.05, *n* = 640 for each instruction, Welch’s *t* test; Fig. 3a; see Supplementary Fig. 4 for changes in the high-γ features of EA02 over days). These differences are attributed to both the subject’s intention to present the images of instructed meaning and the subject’s perception of the different images actually shown as feedback images during the trials. To assess cortical activity without the effects of perception, EC01 and EA02 performed another session of the online task but with their eyes closed (open-loop condition). During the session, an experimenter read out the instructions, which were the same as those in the session for the closed-loop condition, shown on the screen to the subjects, and the subjects, whose eyes were closed, tried to control the feedback images via the same strategy as in the closed-loop condition. In the open-loop condition for EC01, the feedback images rarely represented the instructed meanings (“human” or “tool”)

**Fig. 4.**
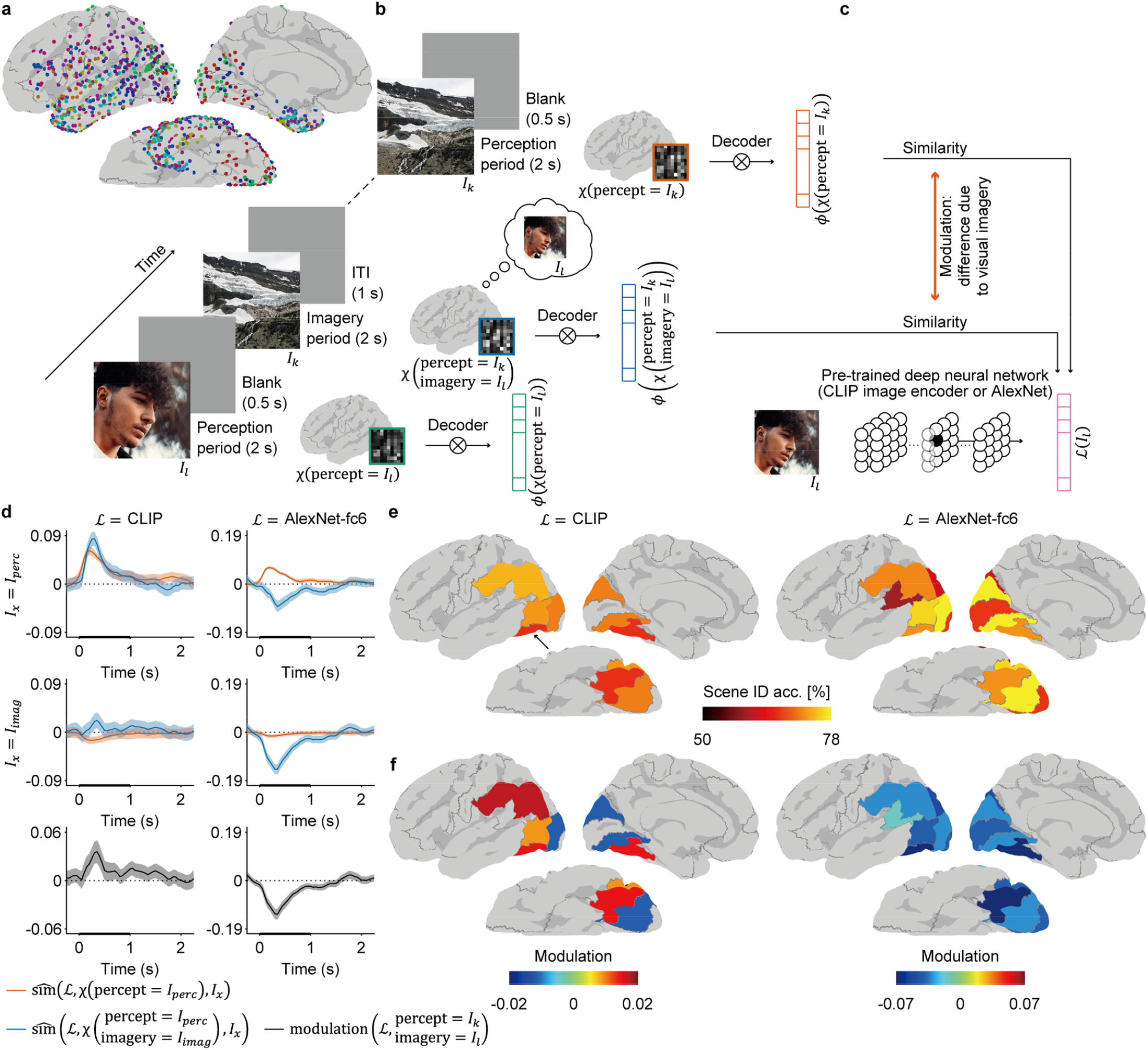
Modulation analysis. (a) For subjects who participated in the modulation task, the locations of individual subdural electrodes were mapped onto the left hemisphere of the normalized brain surface with the colour-coding of the subject. (b) Each trial of the modulation task consisted of a perception period (2 s), a blank period (0.5 s), and an imagery period (2 s). During the perception period, the subjects simply perceived the presented image (*I*); during the imagery period, the subjects perceived another image while visually imagining the image presented during the perception period. Using the decoder (*ϕ*) trained from the image perception task, the brain activity (*χ*) was decoded to infer a vector (*ϕ*(*χ*)). (c) The effect of visual imagery on the inferred vector was assessed by comparing the brain activity while simply viewing an image (*I*_*k*_) (*χ*(percept = *I*_k_)) to that while viewing the same image while visually imagining another Image 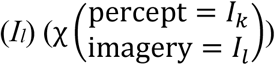. The relative similarity 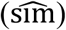 of the inferred vectors from both brain activities to the latent vector of the imagined image (ℒ(*I*_*l*_)) was evaluated 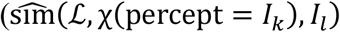 and 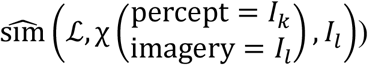to reveal the difference (modulation). (d) The relative similarities during the perception and imagery periods (blue and red lines, respectively) and modulation (black line) calculated for the ventral stream cortex (shown by a black arrow in (e)) are visualized with their corresponding 95% CIs. The stimulus is presented at time 0. (e) Significant scene-identification accuracies were colour-coded on the normalized brain surface for both the CLIP and AlexNet-fc6 latent spaces (FDR-adjusted *p* < 0.05, one-sided paired *t* test against chance level estimated by permutation). (f) Modulation between 0 and 1 s (shown as a thick horizontal axis in (d)) was averaged to be shown with colour-coding on the normalized brain for significant brain regions (FDR-adjusted *p* < 0.05, two-sided one-sample *t* test).

(Supplementary Fig. 2c and d). However, the inferred CLIP vectors were different among trials of the two instructions, resulting in an online accuracy of 72.50%, which was significantly higher than chance level (*p* = 0.003, *n* = 40; retrieval accuracy: 30.42%, *p* = 1.000). In addition, the high-γ features significantly differed between the two instructions in the visual area (FDR-adjusted *p* < 0.05, *n* = 640 for each instruction; Fig. 3b), suggesting that the imagery of the instructed meaning induced significantly different high-γ activity that made the inferred vector closer to the latent vector of the target instruction compared to that of the nontarget instruction for EC01. For EA02, most of the feedback images represented tool images, and the accuracy was at chance level (50.00%, *p* = 0.563, *n* = 40; retrieval accuracy: 52.92%, *p* = 0.204; Supplementary Fig. 2e and f); nevertheless, the high-γ features were significantly different between the two instructions in the visual area, although the differences were smaller than in the closed-loop condition (FDR-adjusted *p* < 0.05, *n* = 640 for each instruction; Fig. 3c and d), suggesting that the imagery induced significantly different high-γ activity even with poor changes in the feedback images in the open-loop condition for EA02. Therefore, it was suggested that the imagery contributed to changes in the inferred vector and that the closed-loop condition helped the control of the inferred vector closer to the instructed meanings.

### Retrieval with the AlexNet-fc6 vector

EA02 and two other subjects (EB01 and EB02) performed the same online task with the images encoded in the AlexNet-fc6 latent space. Here, the decoder was also trained with the high-γ features from the image perception task. When the performance of the decoder was evaluated with the test images in the image perception task, all three subjects showed significant scene-identification accuracy (Table 1); notably, EA02 (and EB02) showed higher scene-identification accuracy when using AlexNet-fc6 vectors than when using CLIP vectors. However, when EA02 performed the online task with the AlexNet-fc6 latent space, the online accuracy decreased to 50.00% (*p* = 0.563, *n* = 40; Table 1 and Supplementary Fig. 5a and b), even though the same electrodes, feedback image pool, instructions, and method for calculating decoding features that showed significant online accuracy with the CLIP vectors were used. Similarly, EB01 and EB02 were not able to control the inferred AlexNet-fc6 vector to be closer to the instructed vector (EB01, 52.50%, *p* = 0.437, *n* = 40; “animal” and “landscape”; EB02, 57.50%, *p* = 0.215, *n* = 40; “animal” and “vehicle”; Table 1 and Supplementary Fig. 5c–f; also see Methods for the detailed conditions of the online task). Notably, the image pool used to retrieve feedback images for these two subjects was the 1,200 training images used in the image perception task; thus, even though the decoder was familiar with the feedback images, the subjects failed to retrieve feedback images with instructed meaning. Although it is difficult to exclude differences in training effects, electrode location, and feedback images, these results suggest that the choice of the latent space affects the controllability of the feedback image in the closed-loop condition, regardless of the high accuracy of the decoders in inferring the latent vectors corresponding to the perceived images.

### Affect of imagery on the inferred vector

The closed-loop online task required the subjects to keep viewing feedback images while imagining the images of instructed meaning so that the feedback images represented instructed meaning. By imagining the instructed meaning while the subjects perceived the feedback images, which were not always the images of the instruction, the subjects succeeded in making the inferred vector closer to the instructed vector with the CLIP latent space but failed with the AlexNet-fc6 latent space. To reveal how the choice of latent space for neural decoding affects the changes in the inferred vectors by visual imagery while the subject perceives images, a modulation task was performed after the image perception task for nine subjects, who were implanted with subdural electrodes around the temporal, occipital, and parietal cortices (Fig. 4a). In the modulation task, each trial started with a 2-s presentation of an image for the subjects to memorize (perception period), followed by a 0.5-s blank screen, after which another image was presented to the subject for 2 s (imagery period) (Fig. 4b). During the imagery period, the subject was instructed to imagine the memorized image visually. The images for the perception and imagery periods were selected from different categories (five images for each category: “face”, “landscape”, “word”, “animal”, “food”, “human”, “scenery”, “plant”, “tool”, and “vehicle”; see Methods and Supplementary Fig. 6).

To analyse the regional differences in neural decoding and the effects of imagery, the ECoG signals of all nine subjects were merged to create a virtual subject (Fig. 4a). For each cortical region (HCP percellation^26^), a decoder (*ϕ*) was trained to infer latent vectors via the high-γ features of the ECoG signals during the image perception task. The decoder was applied to the ECoG signals (χ (·)) of the modulation task to infer a vector while perceiving a *k*-th image (*I*_k_) (*ϕ*(χ (percept = *I*_k_))) and a vector while perceiving a *k*-th image with visual imagery of an *l*-th image 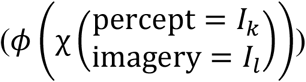 Fig. 4b). The closeness (similarity) of these vectors imagery = *I*_l_ against the latent vector of an *m*-th image (ℒ(*I*_*m*_)) was evaluated via a cosine similarity imagery = *I* metric (sim (*ϕ*(*χ*(percept = *I*_k_)), ℒ(*I*_*m*_)), or sim 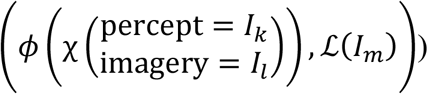 (Fig. 4c). To measure similarity relative to all other images shown in the modulation task, relative similarity was defined as the similarity subtracted by average similarity against latent vectors of all images shown to the subjects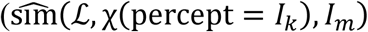, or 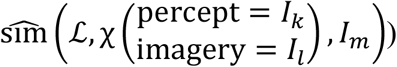 (see Methods).

The time course of the relative similarity against perceived images and imagined images in the ventral stream visual cortex are shown in Fig. 4d (the horizontal axis shows the time in the middle of the 500-ms time window; for other cortical regions, see Supplementary Fig. 7). The relative similarity during the perception period (red lines) against the perceived images (top row) became positive after the image was presented for both the CLIP and AlexNet-fc6 latent spaces, suggesting that the decoders successfully inferred the perceived images. On the other hand, interestingly, the relative similarity during the imagery period (blue lines) against the imagined image (centre row) became positive and negative for the CLIP and AlexNet-fc6 latent spaces, respectively; by subtracting the relative similarity to the same (imagined) images during the perception period as a baseline (red lines in the centre row), the effect of the imagery on the inferred vector was evaluated as modulation (bottom row; modulation 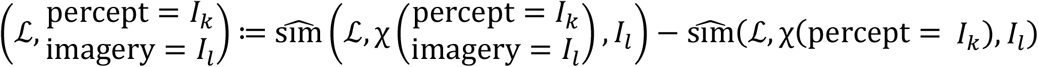; see Fig. 4c). The modulation still became positive for the CLIP latent space and negative for the AlexNet-fc6 latent space. Hence, visual imagery affects the inferred vectors in the ventral stream visual cortex in the CLIP and AlexNet-fc6 latent spaces in the opposite direction.

Modulations were evaluated for all cortical regions that showed significant scene-identification accuracy with the test images in the image perception task (FDR-adjusted *p* < 0.05, one-sided paired *t* test against chance level estimated by permutation; Fig. 4e) because the modulation depends on the decoder. The modulation averaged from 0 to 1 sec after presentation of the visual stimuli is shown in Fig. 4f only for significant modulations (FDR-adjusted *p* < 0.05, two-sided one-sample *t* test against no modulation (0)). When the ECoG signals of higher visual areas were used, the inferred CLIP vector became significantly closer to the latent vector of imagined meaning by the imagery; in contrast, the AlexNet-fc6 vector inferred from the same ECoG signals became significantly away from the latent vector of imagined meaning. Although it is difficult to directly compare the modulations between the two latent spaces, differences in the signs of the modulations demonstrate that the latent space used to represent the visual-semantic information of the images significantly affects the modulation of the inferred vector by the imagery.

## Discussion

In this study, we demonstrated that image retrieval based on CLIP vectors decoded from ECoG signals of the visual area allows the subjects to retrieve images representing the intended meaning from 2.3 million images in a closed-loop condition with a two-choice accuracy of over 90%.

Several factors contributing to the successful retrieval of images via BCIs have been identified. (1) Compared with the open-loop condition, the closed-loop condition improves the online performance. At the beginning of each trial of the close-loop online task, the subjects imagined images while perceiving the blank (grey) screen instead of the feedback images. In the absence of feedback, the similarity to the target instruction vector was often lower than the similarity to the non-target instruction vector. This is in line with previous studies that demonstrated low accuracy in identifying imagined objects (without feedback) using a decoder of perceived visual stimuli^9,15^. However, as feedback continued, similar to our previous study^17^, the similarity to the target instruction vector became higher than the similarity to the nontarget instruction vector (Fig. 2c and f). In addition, online accuracy was greater in the closed-loop condition than in the open-loop condition. Therefore, it was suggested that closed-loop conditions improve online performance. (2) There was a training effect to retrieve images in the closed-loop condition. Multiple days of training to use the BCI improved both the online and retrieval accuracies for one subject (Table 1). (3) Online performance is attributed to changes in the inferred visual–semantic vectors caused by attention and imagery. Previous studies have demonstrated that a superimposed image can be changed to an image that a subject attends to^18,21^, revealing that attention can modulate the output of visual neural decoding. On the other hand, the significant online accuracy of EC01 in the open-loop condition suggested that visual imagery also contributed to the changes in the inferred vectors. (4) More interestingly, it was suggested that the controllability of the feedback image depends on the selection of the latent space to encode the images. The use of the CLIP and AlexNet-fc6 latent spaces showed large differences in closed-loop online performance and modulation of inferred vectors by the imagery, although both latent spaces showed similar (zero-shot) decoding accuracies to infer unseen visual stimuli (Table 1). The modulation was significantly positive when the CLIP latent space was used but significantly negative when the Alexnet-fc6 latent space was used in the inferior parietal cortex, MT+ complex & neighbouring visual areas, and ventral stream visual cortex (Fig. 4f and Supplementary Fig. 7), even though the modulation was assessed for the same ECoG signals. Although it is difficult to directly compare the strength of modulations between latent spaces, the significantly positive and negative modulations cause opposite effects in the inferred vector by the imagery. These results suggest that the appropriate latent space allows significant modulation by imagery and successful control in the closed-loop condition.

The neural information captured by the decoder depends on the latent space^27,28^ so that the online performance changes depending on the properties of the latent space. During the online task using the AlexNet-fc6 latent space, the decoded images seemed to be affected by the attended visual stimuli rather than the imagery; EA02 reported that looking at the corner of the feedback image made the feedback images geometric patterns regardless of the imagery. On the other hand, three subjects, including the above subject (EA02), succeeded in closed-loop control of the decoded image by imagery via the CLIP latent space, suggesting that the use of the CLIP latent space enabled the neural decoding of semantic representations that could be manipulated by imagery (at least partially) independently of perception. These differences might be attributed to the properties of the latent space representing visual– semantic information^29,30^. CLIP consists of a coupled image encoder and text encoder that are trained simultaneously so that the outputs of both encoders acquire a common representation; this might explain the higher neural decoding accuracy in the higher visual areas, which correspond to the areas of common representation of vision and language^31^, than in the early visual areas when decoding was performed in the CLIP latent space (Fig. 4e). In addition, the higher visual area is known to have greater similarity of neural activity during perception and imagery than the early visual areas^32,33^. Consistent with this finding, the modulation using the CLIP latent space was high in the same area (Fig. 4f). However, the hierarchical structure of visual information and its correlation with the latent space are controversial^34-36^. Notably, the correlation varies significantly depending not only on the model architectures and training methods^37^ but also on the training data^38,39^. Indeed, the largest modulation among the visual areas was observed in V1 when the CLIP image encoder model (vision transformer) was trained on only 1,000 categories of images used to train the AlexNet model (Supplementary Fig. 8), although the decoding accuracy using the CLIP latent space was not significant in V1 (Fig. 4e) and the modulation could not be calculated (Fig. 4f). These results demonstrated that the decoded information differs depending on the latent space even for the same neural activities. A comparison of decoded information using different latent spaces may reveal neural representations during perception and imagery.

In this study, the small number of subjects with different conditions might limit the generalizability of the results. In particular, electrode location (Supplementary Fig. 1a) seems to be one factor that affects online performance. Moreover, individual differences in the vividness of imagery^40,41^ and strategies to imagine^42^ might affect online performance. Further studies with more subjects and various electrode locations will reveal the influence of cortical area and individual characteristics on online performance in detail. Nevertheless, some subjects who performed the closed-loop online task with multiple latent spaces still demonstrated how the selection of the latent space affects the online performance (EA02 for CLIP and AlexNet-fc6; see Supplementary Fig. 9 for AlexNet-fc6 and word2vec). In addition, modulation analysis was performed on the same ECoG signals for both the CLIP and AlexNet-fc6 latent spaces, regardless of the difference in the individual electrode locations; the results still suggested that the differences in latent space cause differences in modulation.

## Conclusions

Visual–semantic neural decoding using an appropriate latent space in a closed-loop condition enables a BCI to retrieve images from a large image dataset via visual cortical ECoG signals.

## Methods

### Subjects

For this study, eleven subjects with drug-resistant epilepsy (six males; 26.7 ± 11.0 years old, mean ± standard deviation (SD); Supplementary Table 1) were obtained from three university sites (Osaka University, Juntendo University, and Nara Medical University). The intracranial electrodes were implanted in these subjects at each university hospital for the purpose of treating epilepsy (number of subdural electrodes: 64.9 ± 19.4; number of depth electrodes: 6.2 ± 9.3; Supplementary Fig. 1a and b). Before participation in the study, written consent was obtained from each subject after the nature and possible consequences of the study were explained. The experiment was performed in accordance with the experimental protocol approved by the ethics committee of each hospital (Osaka University Medical Hospital: Approval No. 14353, UMIN000017900; Juntendo University Hospital: Approval No. 18-164; Nara Medical University Hospital: Approval No. 2098).

### Sample size

The amount of data collected per subject was determined by the clinical treatment schedule and the amount of time each subject was willing to volunteer for this study. The number of trials in each task was determined on the basis of our previous study^17^. The reproducibility of the online task was validated by independent multiple subjects.

### Localization of intracranial electrodes

The intracranial electrodes were located on presurgical T1-weighted magnetic resonance (MR) images and postsurgical computed tomography (CT) images via the following procedure. (1) The individual cortical surface was extracted and registered to a template brain (fsaverage) from the MR images via FreeSurfer^43^. (2) Using the BioImage Suite^44^, each intracranial electrode was manually located on the CT images, which were coregistered to the MR images. (3) The located subdural electrodes were projected onto the individual cortical surface via the intracranial electrode visualization toolbox^45^. (4) Using the registration in (1), the location of each subdural electrode on the template brain was determined. (5) For further region-based analysis, the subdural electrodes were assigned to one of 22 regions on the basis of the parcellation of the human connectome project (vid. supplementary neuroanatomical results)^26^.

### Image datasets

In this experiment, the following six image datasets were used as visual stimuli in the image presentation task and modulation task, and as feedback images for the online task. (1) The GOD image dataset^9^ consisted of 1,200 training images from 150 categories (8 images per category) and 50 test images from 50 categories (1 image per category). (2) The preceding stimuli image dataset consisted of 10 images from 10 categories (1 image per category). (3) The baseline image dataset comprises sixty images. (4) The three-category image dataset contained five images each for the “face”, “landscape”, and “word” categories (Supplementary Fig. 6a). The images originated from still images extracted from stimuli movies used in our previous study^17^. (5) The seven-category image dataset comprises five images for each category: “animal”, “food”, “human”, “scenery”, “plant”, “tool”, and “vehicle” (Supplementary Fig. 6b). (6) The Unsplash image dataset^25^ consisted of 2,283,429 high-quality images. For the creation of these datasets, all images were preprocessed by cropping them into squares following the methods of a previous study^9^. The images of (1) the GOD image dataset, (2) the preceding stimuli image dataset, (3) the baseline image dataset, and (5) the seven-category image dataset originated from ImageNet^24^ and included no overlapping images. Although other datasets originated from different image sources, the overlap of images between the datasets was confirmed via AlexNet-fc6 vectors of all images in the datasets; the maximum cosine similarity was 0.8680 between the ImageNet-originated datasets and the three-category image dataset, 0.9679 between the ImageNet-originated datasets and the Unsplash image dataset, and 0.9620 between the three-category image dataset and the Unsplash image dataset, indicating that there was no overlap among the images.

### Extraction of latent vectors

Images from the image datasets were encoded into latent vectors via pretrained CLIP and AlexNet models. For the extraction of the CLIP vectors, a custom program based on PyTorch 20.03 was used to acquire unit responses at the output of the pretrained CLIP image encoder model (ViT-B/32)^22^. On the other hand, for the extraction of the AlexNet-fc6 vectors, a custom program based on Chainer 4.5.0 was used to extract the unit responses from the fc6 layer of the pretrained BAIR Reference CaffeNet model^46^. To reduce the computation time, 1,000 units were randomly selected from 4,096 units; the selection was fixed throughout this study.

### ECoG acquisition

During the experiments, the subjects either sat on beds in their hospital rooms or were seated on chairs in front of a computer screen to view the visual stimuli or real-time feedback images. The ECoG signals from each subject were recorded by an EEG-1200 (Nihon Koden, Tokyo, Japan) at 10 kHz with reference to the average of two intracranial electrodes. The presentation timing of the visual stimuli and real-time feedback images were monitored by DATAPixx3 (VPixx Technologies, Quebec, Canada) and recorded synchronously with the ECoG signals.

### Experimental procedures

All eleven subjects participated in the *image presentation task* to train and evaluate a decoder (Supplementary Fig. 1c and d, and Supplementary Table 1). With the decoder trained by the image presentation task, nine subjects participated in the *modulation task* to evaluate changes in cortical activity via visual imagery (Fig. 4b), whereas five subjects participated in the *online task* (Fig. 1b and Supplementary Fig. 1e). Moreover, to compensate for the change in electrode impedance among the recording sessions, a *baseline recording task* was performed at the beginning of each recording session. For all tasks, the subjects were required to keep their eyes on a fixation point at the centre of the screen.

#### Baseline recording task

To compensate for the recording-session-to-recording-session difference in electrode impedance, the baseline recording task was performed at the beginning of each recording session. The task consisted of one run (one session), in which the subjects were presented with images from the baseline image dataset in random order without a blank screen between the presentations. The duration of the presentation of each image was 1125 ± 25 ms.

#### Image presentation task

All the subjects participated in the image presentation task, in which images from the GOD image dataset were presented as visual stimuli. One training session consisted of two runs to present all the GOD training images, where one test session consisted of one run. In each run, 10 images of the preceding stimuli image dataset were first presented in random order, followed by images from the GOD image dataset in random order. There were no blanks between the images. The duration of the presentation of each image was 525 ± 25 ms.

#### Modulation task

To reveal changes in cortical activity due to visual imagery, nine subjects participated in the modulation task once for both the three-category image dataset (three runs) and the seven-category image dataset (five runs). For each trial, the image to be visually imagined during the imagery period was first presented for 2 s (perception period), followed by a 0.5-s blank and presentation of another image for 2 s (imagery period) (Fig. 4b). The subjects were instructed to memorize the image presented during the perception period and to visually imagine the memorized image in the following imagery period. The intertrial interval (ITI) was 1 s. The set of images to be presented during the perception period and the imagery period was determined as follows. (1) For the trials of the three-category image dataset, all possible pairs between the images from another category were used, resulting in 150 pairs (trials). (2) For the trials of the seven-category image dataset, five groups of images without overlap among the groups were first created by selecting one image from each category; within each group, all possible pairs were created, resulting in 240 pairs (trials). The grouping was kept consistent throughout this study. The order of the trials for the three-category and seven-category image datasets were randomized and divided into three or five runs, respectively.

#### Online task

Five subjects participated in the online task to evaluate the online performance of the decoder trained from ECoG signals during the image perception task (for detailed conditions, see Table 1). To determine the set of target instructions, the decoder was set in free-running mode, showing feedback images every 250 ms. During this period, the experimenters showed some representative images of candidate categories (Supplementary Fig. 6) and asked the subjects if they could control the feedback images to show the images of the same category. Notably, the subjects were not instructed to show the same images but rather the images of the same category during the following online task. Each trial started with the presentation of a target instruction visually (EC01, EA01, and EA02) or auditorily (EB01 and EB02) in Japanese, followed by a 2.5-s or 1-s blank screen, respectively, and 32 feedback images, each with a duration of 250 ms (e.g. Fig. 2b). The subjects were instructed to control these feedback images to show the category of the target instruction by using visual imagery. In the session for EA02 on Day 6, success/failure of the trial was evaluated on the basis of the cosine similarity and was presented for 2 s after the presentation of the feedback images.

### Signal preprocessing

Noise-free channels were first identified by visual inspection for exclusion from further analyses. For the online task, channels considered in regions unrelated to the control (e.g., motor area) were also excluded. The ECoG signals recorded for each task were re-referenced by common averaging across the remaining channels.

### Calculation of high-γ features

Given the 500-ms preprocessed ECoG signals of a channel (*X*_*signal*_(*t*)), the signals were applied with a Hamming window and fast Fourier transformation (FFT) to acquire either the amplitude spectrum density (*ASD* (*X*_*signal*_(*t*)) or the power spectrum density (*PSD* (*X*_*signal*_(*t*))). The amplitude/power spectrum density was then averaged within the high-γ frequency band to acquire the raw amplitude/power spectrum density (asd/psd) features as follows.

asd features: 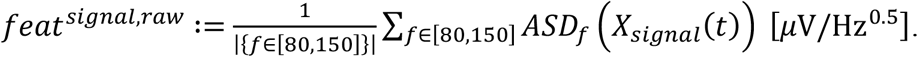

psd features: 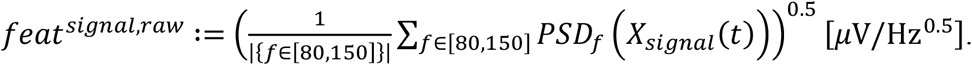

The raw asd/psd features (*feat*^*signal,raw*^) were then compensated (standardized) for the recoding-session-to-recording-session change in the electrode impedance using the ECoG signals measured during the baseline recording session performed at the beginning of each recording session. Baseline features were obtained from the 500-ms ECoG signals during presentation of the images from the baseline image 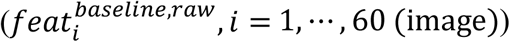.By comparing these baseline features against another set of baseline features in the baseline recording session just before the first run of the training session of the image presentation task 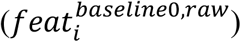, the compensation factor was calculated as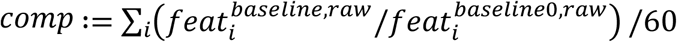. The compensation factor was then applied to the raw features as follows to acquire the high-γ features: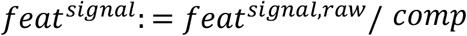

### Construction of the decoder

Throughout this study, a ridge regression model was used as a decoder to infer the latent vectors from the high-γ features. The best parameter for the model was estimated via 10-fold cross-validation^47^. For each fold, the high-γ features in the training samples of the fold were standardized by *z*-scoring using the mean and SD for each dimension of the features; the same mean and SD were also applied to the high-γ features in the testing samples of the fold to avoid data leakage^48^. For each of the regularization parameters (10^−8^, 10^−7^, ⋯, 10^8^), a regression model was trained and applied to the testing samples. To select the optimal regularization parameter, the dimensionwise correlation between the true and inferred latent vectors of all samples was calculated for each dimension of the latent space and averaged among the dimensions; the parameter with the highest average was selected. The final decoder model was trained with all the high-γ features and the selected regularization parameter. Notably, the regression models used in this study did not have constant terms because, empirically, the models with the constant term deteriorated scene-identification accuracies in some latent spaces.

### Real-time decoding in the online task

#### Decoder training

For each subject, a decoder model for inferring the latent vector was trained (Fig. 1a) using all high-γ asd/psd features acquired during the presentation of the GOD training images (0– 500 ms). For cross-validation during decoder training, the trials of the image presentation task were split by selecting five consecutive trials.

#### Real-time feedback

In the online task, the ECoG signals were acquired and decoded in real time to infer a latent vector and select a feedback image on the basis of the inferred vector. The ECoGs of the most recent 500 ms (*χ*_*online*_) were re-referenced by preprocessing, converted into raw features, compensated for recording-session-to-recording-session impedance changes of the electrodes, and applied with the decoder model (*ϕ*) to infer the latent vector (*v*_*online*_ = *ϕ*(*χ*_*online*_)). Notably, *v*_*online*_ was linearly interpolated for EB01 (*α* = 0.5; see Fukuma et al., 2022^17^). From an image pool 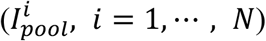, the feedback image was selected on the basis of the highest cosine similarity (or Pearson’s correlation coefficient for EB01 and EB02) 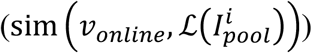 between the inferred vector and the true latent vectors of the images in the pool 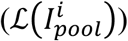; this image search process was performed with the ngt library^49^ to enable fast feedback by the feedback images. The system delay from the acquisition of the ECoG signals to the display of the feedback image was 174.4 ± 29.7 ms (mean ± SD; measured from the closed-loop online task performed by EA02 on Day 1), even for the 2.3 million images in the Unsplash image dataset^25^.

#### Evaluation of the online task

The success/failure of each trial was evaluated on the basis of the similarities between the inferred vectors used to determine the *i*-th feedback images 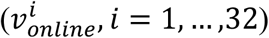 and the latent vectors of all instructions used for the subject. Here, the latent vectors corresponding to instruction 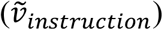 were acquired in the following way. (1) For the CLIP latent space, the CLIP text encoder was applied to the words “animal”, “human”, and “tool”. (2) For the AlexNet-fc6 latent space, images shown to the subject to determine the instructions before the actual trials were used to extract the latent vectors. For each inferred vector, the cosine similarity/correlation (sim) with the latent vector of the target instruction 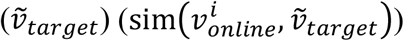 and that of the nontarget instruction 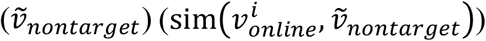 were calculated. These similarities were then averaged among 32 feedback images for each instruction 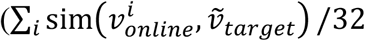and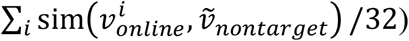. When the average similarity to the target instruction was greater than the similarity to the nontarget instruction 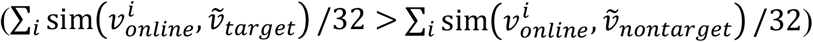,the trial was considered successful.

### Decoding and modulation analysis

#### Decoder training and inference

For the individual decoding analysis, the decoder used in the online task was applied to the high-γ features (0–500 ms) during the perception of the GOD test images. Region-based decoding analyses were conducted by creating a virtual subject who participated in one training and test session of the image presentation task. The high-γ asd features of all the subjects were concatenated across the electrodes to form the decoding features of the virtual subject. In the case of subjects who participated in multiple sessions, the high-γ features were averaged across the trials with the same visual stimuli image. For each cortical region, a decoder was trained with the high-γ (training) features from the subdural electrodes in the region to be applied to the high-γ (testing) features.

#### Scene-identification accuracy

The offline performance of the decoder was evaluated for scene-identification accuracy within the GOD test images. For each test image, the cosine similarity of the inferred vector with latent vectors of other test images was acquired; scene-identification accuracy was defined as the ratio of the image that had a smaller similarity than the similarity between the inferred vector and the corresponding true latent vector^17^. The scene-identification accuracy for each test image was averaged to acquire the average scene-identification accuracy (shown in Table 1 and Fig. 4e).

#### Retrieval accuracy

To assess the controllability of the inferred vector in the latent space, the retrieval accuracy was calculated via seven candidate categories for the online task (Supplementary Fig. 6b), in a similar way to the scene-identification accuracy. For each trial, the average cosine similarity between the 32 inferred vectors used to determine the feedback images and the target instruction vector was compared with the average similarities between the same inferred vectors and latent vectors for the other six categories; the ratio in which the average cosine similarity to the target instruction vector became larger than the other similarities was defined as the retrieval accuracy for the trial.

#### Decoding for modulation analysis

In the modulation analysis, the same decoder used in the region-based decoding analysis (Fig. 4e) was applied to the ECoG signals during the modulation task. For each cortical region, the decoder (*ϕ*) was applied to 500-ms ECoG signals (*χ*_*t*_), whose centre of the time window ranged from 0 ms to 2,000 ms according to the onset of the image presentation with 50 ms intervals (*t* = 0, 0.05, ⋯, 2 [*s*)); hence, an inferred vector was acquired in each trial for both the presentation period of the trial (*ϕ*(*χ*_*t*_(percept = *I*_*k*_))), where the subject perceived an image (*I*_*k*_), and the imagery period of the trial 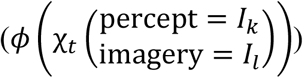, where the subject perceived an image (*I*_*k*_) while imagining an image (*I*_*l*_).

#### Evaluation of modulation

For a given inferred vector (*ϕ*(*χ*_*t*_)) and a given image (*I*), the similarity of the inferred vector to the latent vector of the image (ℒ(*I*)) was defined as follows using cosine similarity (sim): sim(*ϕ*(*χ*_t_), ℒ(*I*)). To evaluate the selectivity of the inferred vector towards the given image among other images presented to the subjects, the relative similarity 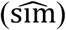 was acquired by subtracting the average similarity between the inferred vector and all images presented to the subject in the task (*I*_*n*_ where *n* =1,…., N) as follows:

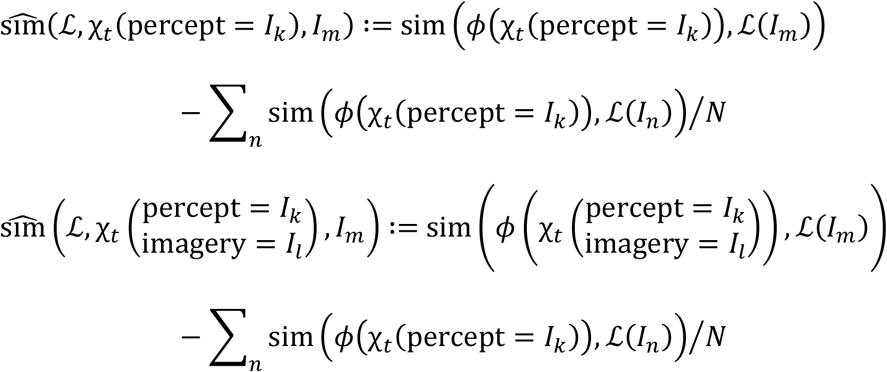

The difference in the relative similarity due to visual imagery (of an image *I*_*l*_ while perceiving an image *I*_*k*_) was defined as the modulation as follows:

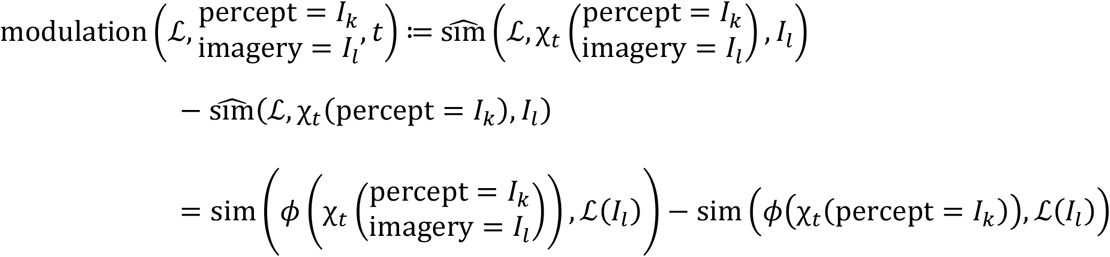

### Illustrations

The images from the GOD image dataset and three-category image dataset shown in this paper were replaced by semantically similar images from the Unsplash image dataset because of the copyrights of the images.

### Statistics and reproducibility

Throughout this study, the α level for the statistics was set to 0.05.

The scene-identification accuracy was tested with a one-sided paired *t* test against the chance level of each GOD test image (Table 1 and Fig. 4e). The chance level for each image was estimated by calculating the scene-identification accuracy by shuffling the correspondence between the inferred vectors and the true latent vectors (permutation). The shuffling was repeatedly performed 1,000 times to be averaged between the repetitions. The acquired *p* values for the cortical regions were adjusted with false discovery rate (FDR) correction (Fig. 4e).

The online accuracy was tested by applying a one-sided binominal test to assess successful control over trials. For EA02, the *p* values were adjusted for multiple comparisons (3 — number of sessions using the same latent space (CLIP)) via Bonferroni correction.

The retrieval accuracy was tested with a one-sided paired *t* test against the chance level of each trial. The chance level was estimated by calculating the retrieval accuracy with randomly selected instructions 1,000 times. The instructions were selected from the seven candidate categories (Supplementary Fig. 6b). For the online task of EA02 using the CLIP latent space, the *p* values were adjusted for multiple comparisons via Bonferroni correction.

The difference in the high-γ features due to the different instructions during the online task was tested with two-sided Welch’s *t* test; the acquired *p* values for all electrodes were adjusted with FDR correction (Fig. 3a–d).

Modulation was first calculated for the cortical regions that showed significant scene-identification accuracy (Fig. 4e). The calculated modulations were then tested with a two-tailed one-sample *t* test against no modulation (0) with FDR correction (Fig. 4f).

## Acknowledgements

We thank all subjects for their participation. This research was supported by the Japan Science and Technology Agency (JST) Moonshot R&D (JPMJMS2012), JST Core Research for Evolutional Science and Technology (CREST) (JPMJCR18A5), JST AIP Acceleration Research (JPMJCR24U2), and Japan Society for the Promotion of Science (JSPS) Grants-in-Aid for Scientific Research (KAKENHI) (JP26560467 and JP20H05705).

## Author contributions

Conceptualization: T.Y.; methodology: R.F. and T.Y.; investigation: R.F., T.Y., H.Sug., K.T., S.O., N.T., Y.I., H.M.K., H.Suz., H.Y., T.I. and M.N.; data curation, formal analysis, and software: R.F.; funding acquisition: R.F., T.Y., and H.K.; writing–original draft: R.F. and T.Y.; writing–review and editing: R.F., T.Y., S.N., Y.K., and H.K.; supervision: T.Y.

