## Supplementary Materials for "Image retrieval based on closed-loop visual–semantic neural decoding"

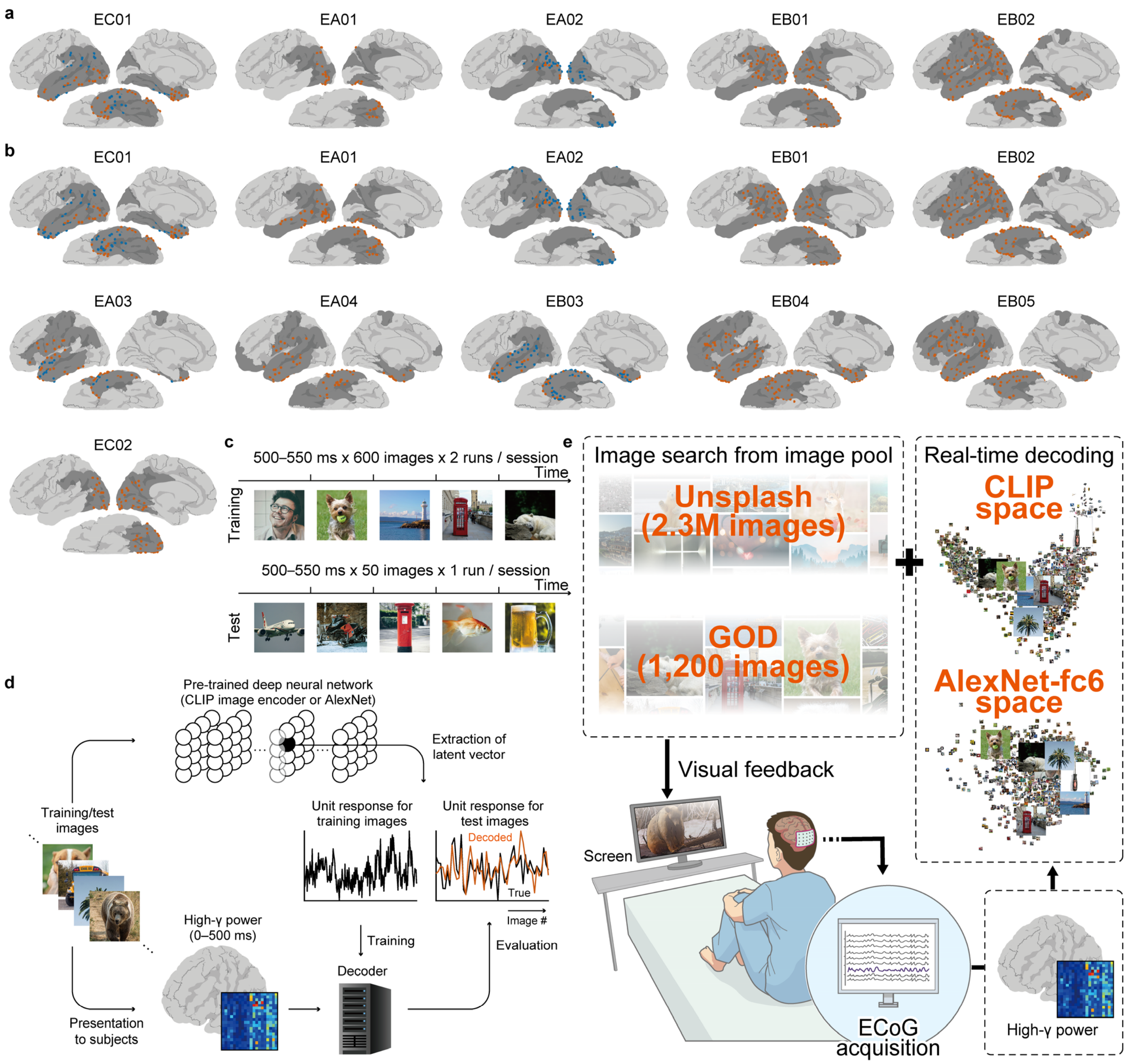


**Supplementary Fig. 1. Electrode location and experimental schematics.**

(a, b) Electrode location of individual subjects used in (a) the online task and (b) the modulation analysis is shown on the normalized brain, with the colour-coding of red and blue denoting the electrodes on the left and right hemispheres, respectively. The cortical area marked with a darker colour denotes the regions where the subdural electrodes were located. (c) The image perception task was composed of a training session and test session, in which the subjects were shown with 1,200 GOD training images and 50 test images, respectively. Each subject participated in one or more training and test sessions while recording ECoG signals (Supplementary Table 1). (d) Images presented to the subject were fed into the CLIP image encoder model or AlexNet model to acquire latent vectors (CLIP vectors or AlexNet-fc6 vectors). A linear decoder model was trained so that these latent vectors of the training images can be inferred from the high-γ features (standardized high-γ powers) during the presentation of the corresponding training images (0–500 ms). To evaluate the performance of the decoder, the decoder was tested on the high-γ features for the test images. (e) Using the decoder acquired in (d), five subjects participated in the online task. From their real-time high-γ powers, a latent vector was inferred in the CLIP or AlexNet-fc6 latent space every 250 ms; the feedback image was searched from the Unsplash image dataset or the GOD training image dataset on the basis of the highest cosine similarity (or Pearson's correlation coefficient for EB01 and EB02) with the inferred vector.

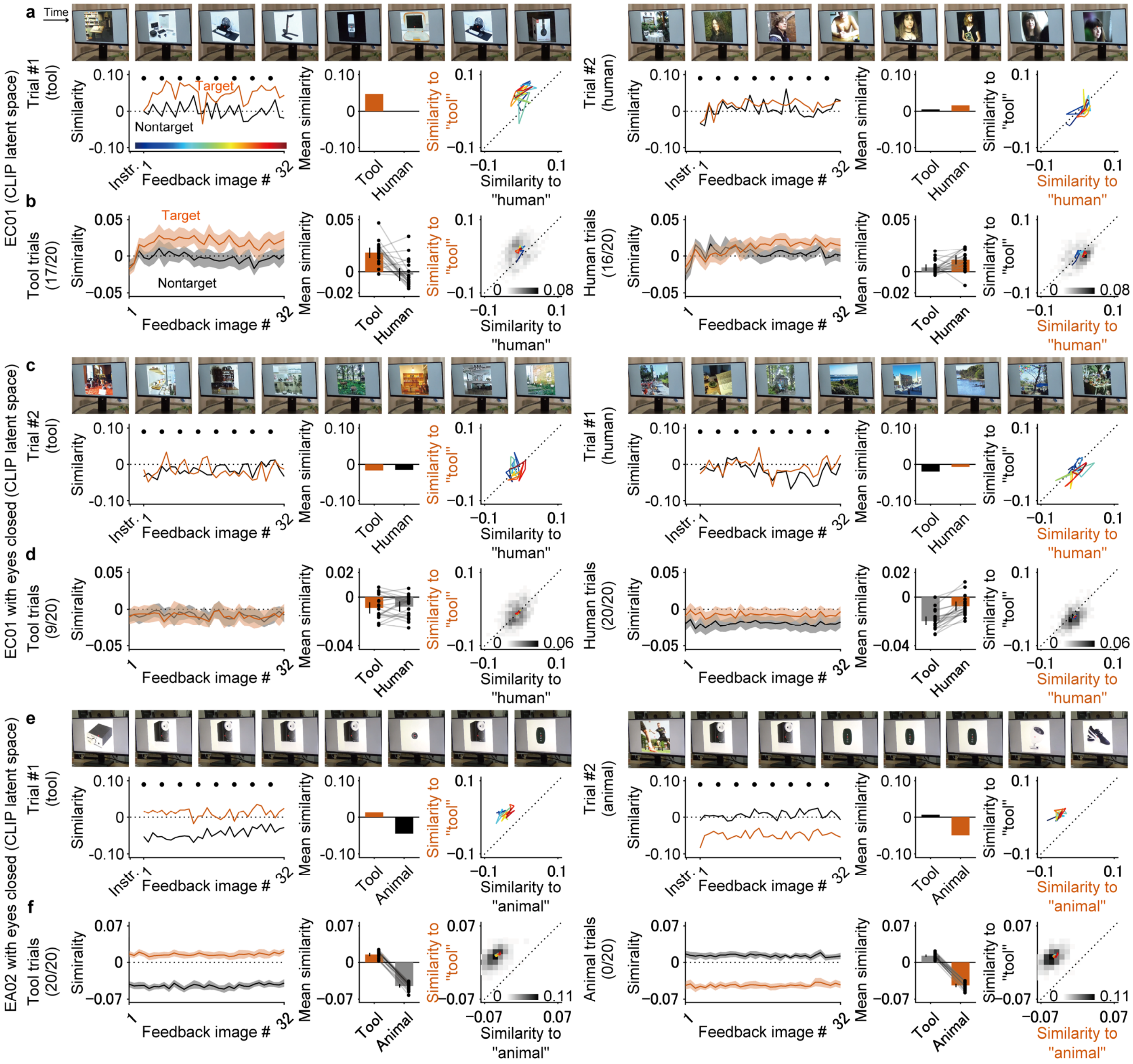


**Supplementary Fig. 2. Online task with CLIP latent space for closed- and open-loop conditions**

(a) Representative trials of EC01 for the online task with a closed-loop condition are shown with pictures of the feedback screen (top panel), cosine similarity of the inferred vector to the target/nontarget instruction vector (left bottom; time course panel), average of the cosine similarity during the trial (centre bottom; mean panel), and trajectory of the similarities during the trial (right bottom; trajectory panel). In the time course panel, the red and black lines denote the cosine similarity of the inferred vector used to search the feedback image with the target and nontarget instruction vectors, respectively. The black dots above the lines denote the time of the pictures shown in the top panel. The mean panel shows the average cosine similarity during the trial, which is shown in the time course panel. By comparing these averages, the success of each trial was evaluated. The time of the trajectory is colour-coded, as shown in the time course panel. (b) Each panel shows the trial average of the cosine similarity in the corresponding panel in (a). The shaded area in the time course panel and error bar in the mean panel denote 95% CIs, where the dots in the mean panel represent the actual value for each trial. In the trajectory panel, the density of the trajectory is shown with colour-coding. (c–f) Representative trials and trial average of the similarities for (c–d) EC01 and (e–f) EA02 using the CLIP latent space with the open-loop condition.

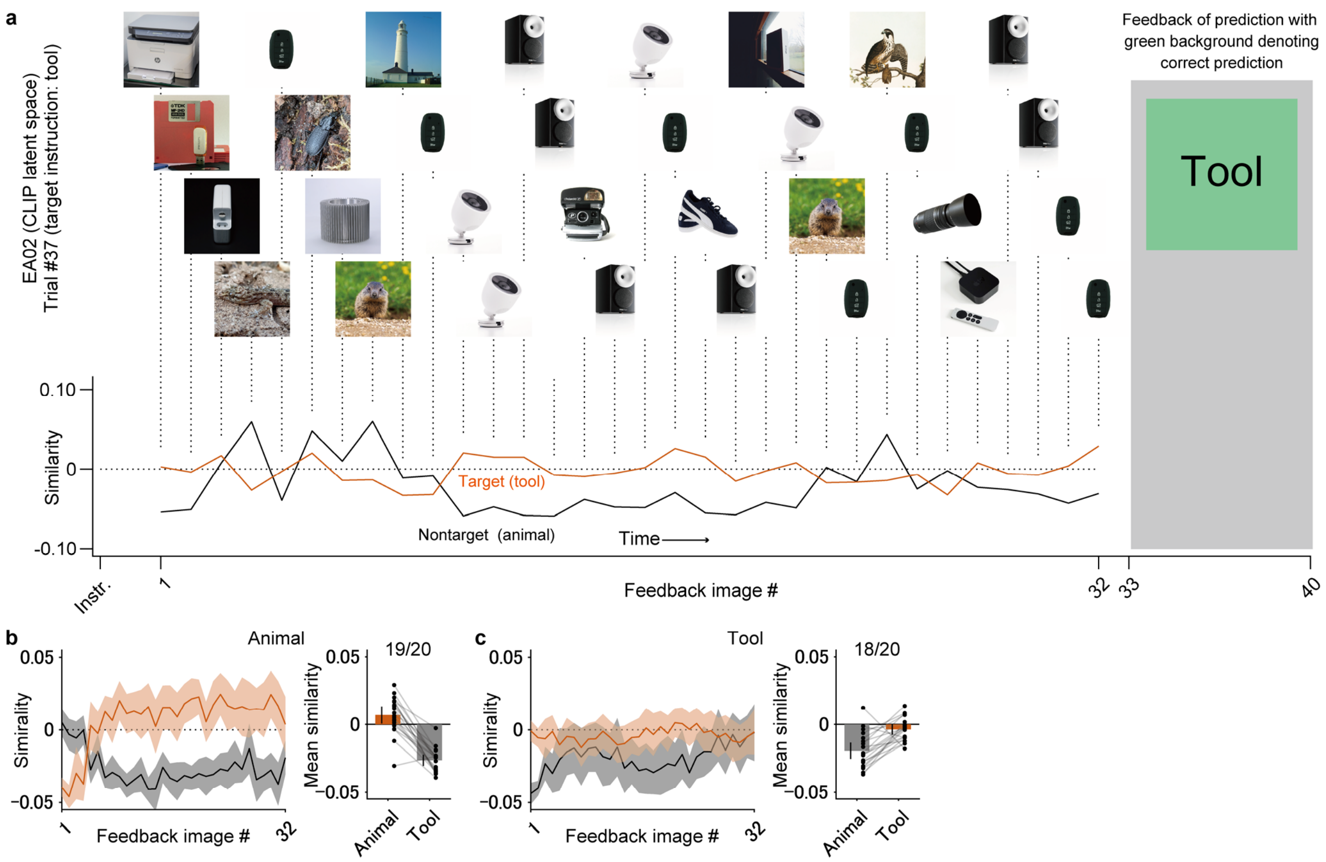


**Supplementary Fig. 3. Closed-loop online task performed by EA02 in the CLIP latent space with success/failure feedback for each trial.**

(a) EA02 performed the closed-loop online task with visual feedback of success/failure at the end of each trial on Day 6. A representative trial (trial #37) from the session of the task is shown with the feedback images (top) and the cosine similarity of the inferred vector to the target/nontarget instruction vector (bottom). At the end of the trial, the inferred category ("animal" or "tool") was displayed with a coloured background. The colours were green and red for correct and incorrect inference, respectively. The duration of the success/failure feedback was 2 s. (b, c) The trial average of the cosine similarity shown in (a) for each order of the feedback image (left) and the trial- and feedback-image-averaged similarity (right) are shown for trials of (b) animal and (c) tool instructions. The shaded area in the left panel and the error bar in the right panel denote 95% CIs.

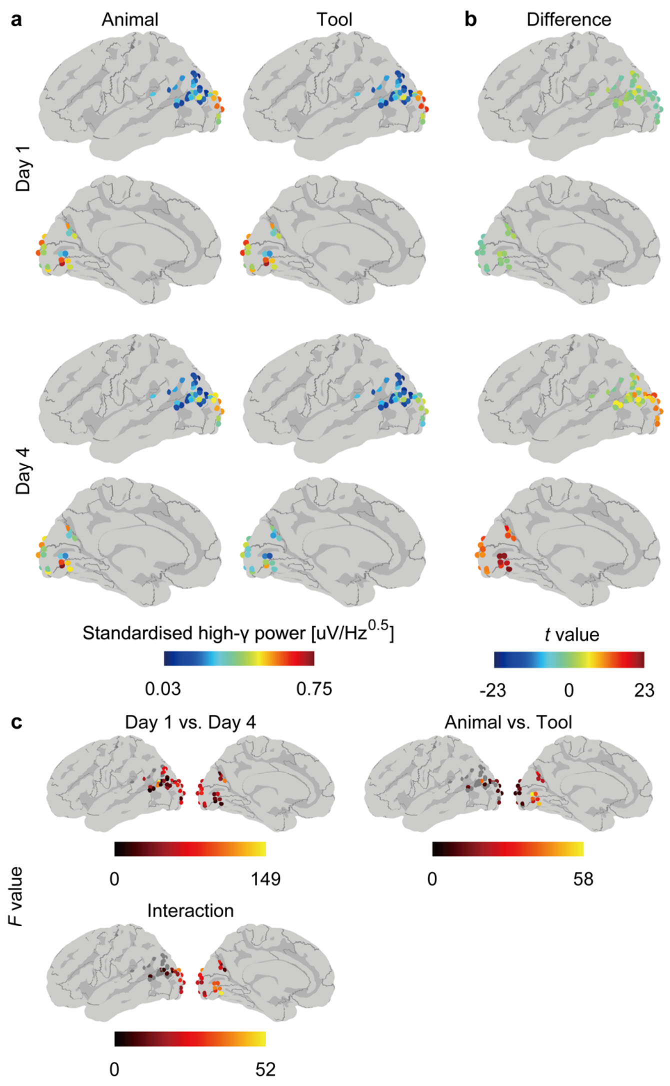


**Supplementary Fig. 4. Cortical activity during closed-loop online task in the CLIP latent space performed by EA02 on Days 1 and 4.**

(a) Trial and feedback image averages of high-γ features (standardized high-γ powers) used to determine the feedback images during the closed-loop online task of EA02 on Days 1 and 4 are visualized on a normalized brain surface with colour-coding. (b) Differences in the high-γ features between the animal and tool instructions (shown in (a)) were evaluated with two-sided Welch's *t* test ($n$ = 640 for each instruction and day) to visualize differences at each electrode. (c) The high-γ features shown in (a) were subjected to two-way ANOVA to visualize significant differences in power due to days, instructions, and interactions (FDR-adjusted $p < 0.05$).

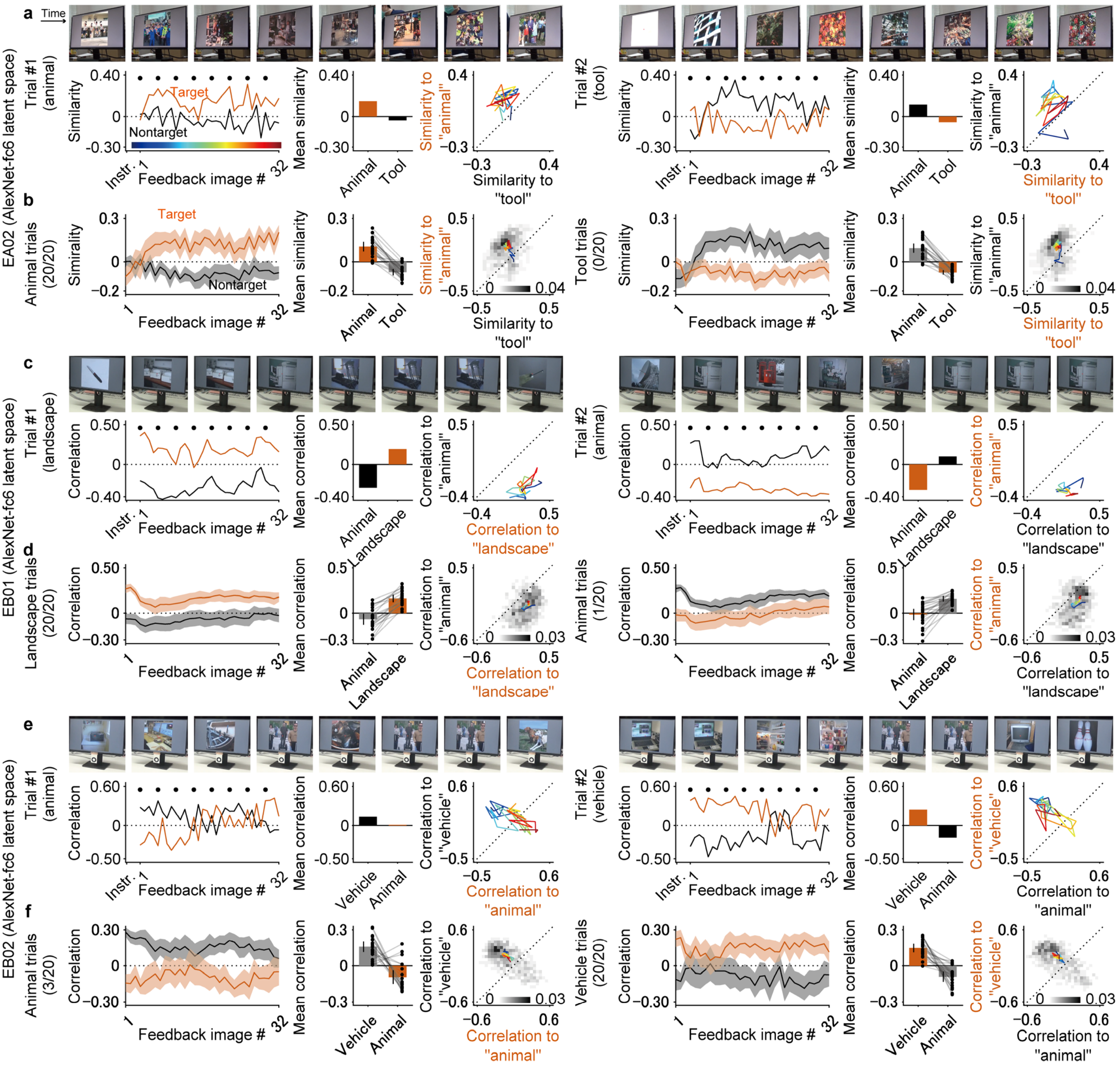


**Supplementary Fig. 5. Closed-loop online task in the AlexNet-fc6 latent space performed by EA02, EB01, and EB02.**

(a) Representative trials for the closed-loop online tasks performed by EA02 with decoding in the AlexNet-fc6 latent space are shown with pictures of the feedback screen (top panel), cosine similarity of the inferred vector to the target/nontarget instruction vector (left bottom; time course panel), average cosine similarity during the trial (centre bottom; mean panel), and trajectory of the similarities during the trial (right bottom; trajectory panel). In the time course panel, the red and black lines denote the cosine similarity of the inferred vector used to search the feedback image with the target and nontarget instruction vectors, respectively. The black dots above the lines denote the time of the pictures shown in the top panel. The mean panel shows the average cosine similarity during the trial, which is shown in the time course panel. By comparing these averages, the success of each trial was evaluated. The time of the trajectory is colour-coded, as shown in the time course panel. (b) Each panel shows the trial average of the cosine similarity in the corresponding panel in (a). The shaded area in the time course panel and error bar in the mean panel denote 95% CIs, where the dots in the mean panel represent the actual value for each trial. In the trajectory panel, the density of the trajectory is shown with colour-coding. (c–f) Representative trials and trial average of the similarities for (c–d) EB01 and (e–f) EB02 using the AlexNet-fc6 latent space in the closed-loop condition.

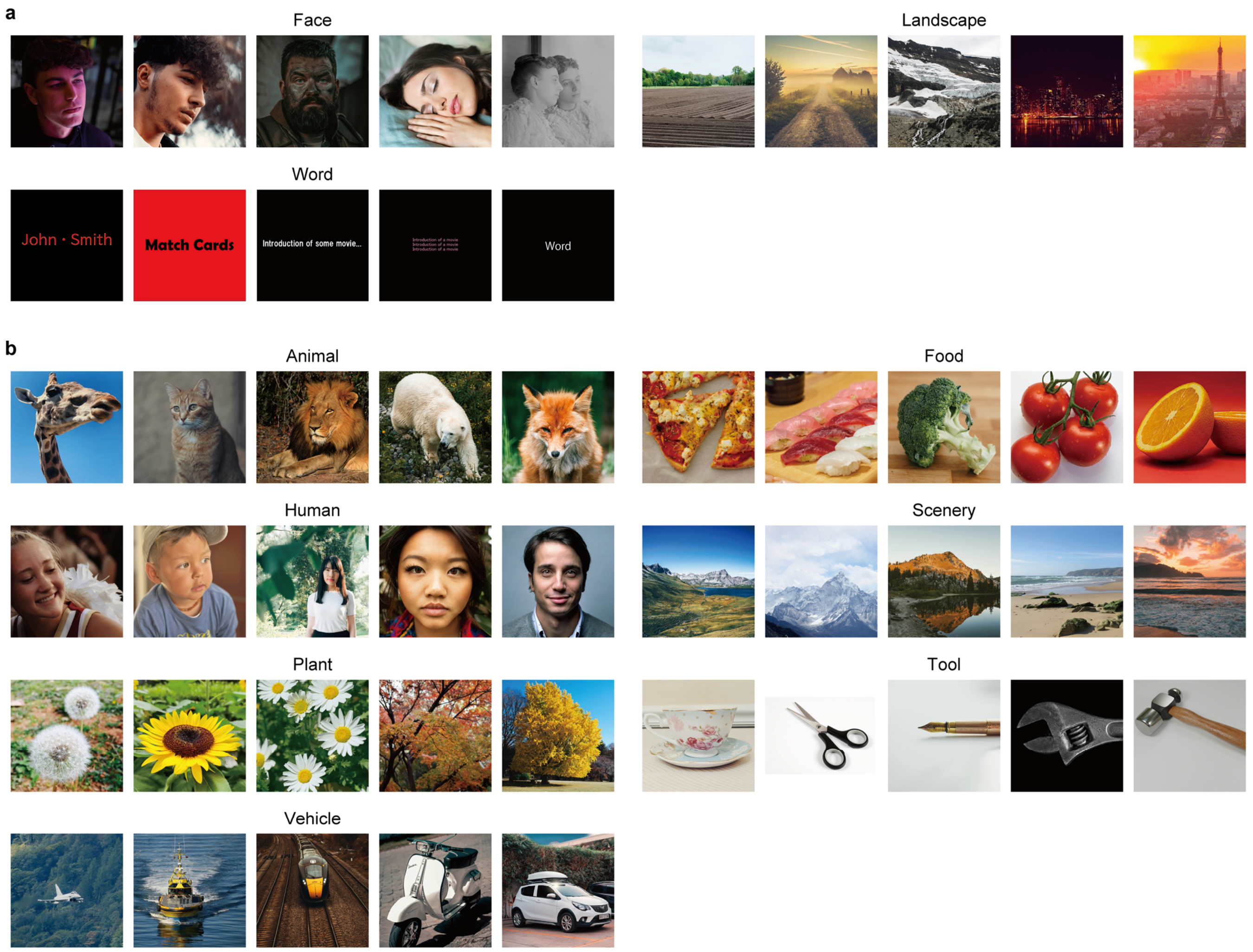


**Supplementary Fig. 6. Images presented during the modulation task.**

(a, b) Images used for the (a) three- and (b) seven-category modulation tasks are shown. Owing to the copyright, semantically similar images from the Unsplash image dataset are represented instead of the actual image.

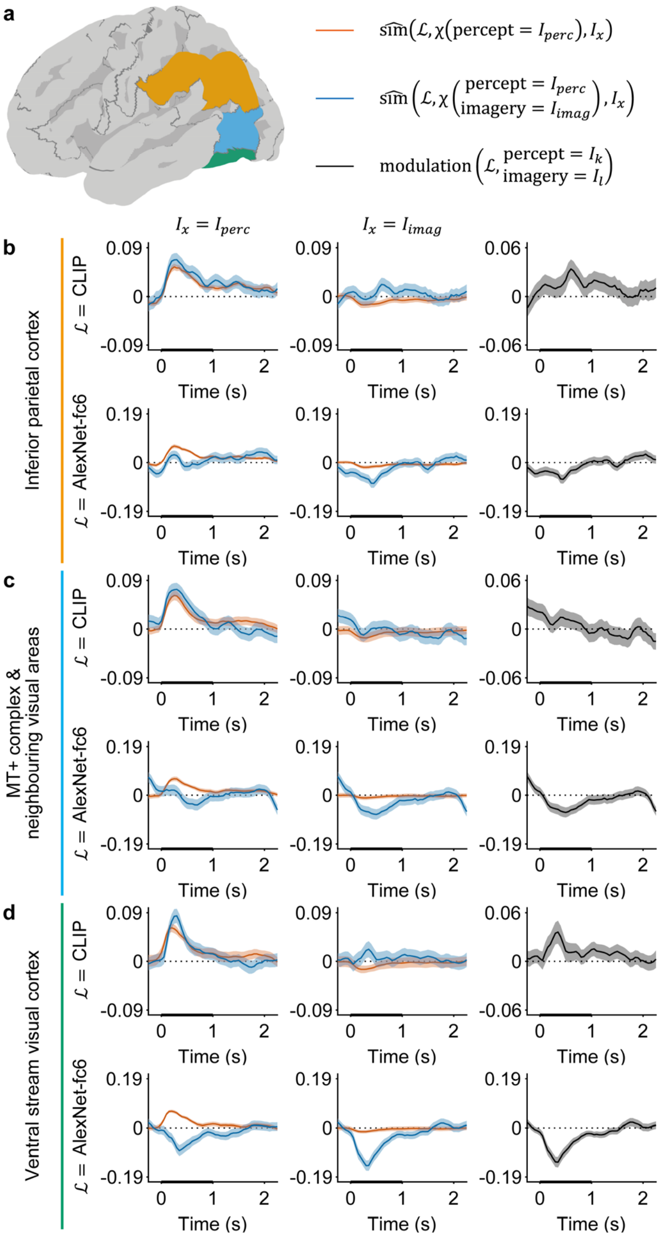


**Supplementary Fig. 7. Time course of modulation in various cortical regions.**

(a) Cortical regions shown in (b–c) are visualized on the normalized brain surface. (b–d) Time course of the relative similarity between the inferred latent vector and latent vector of the presented image (left), that of the relative similarity between the inferred latent vector and latent vector of the imagined image (centre), and that of modulation (right) are visualized for (b) the inferior parietal cortex, (c) the MT+ complex & neighbouring visual areas, and (d) the ventral stream visual cortex. The shaded area denotes the corresponding 95% CIs. The time on the horizontal axis denotes the centre time of the 500-ms time window used for decoding according to the presentation of the visual stimuli (0 ms). The thick horizontal axis from 0 to 1 s denotes the time range in which the average modulation in Fig. 4f was calculated.

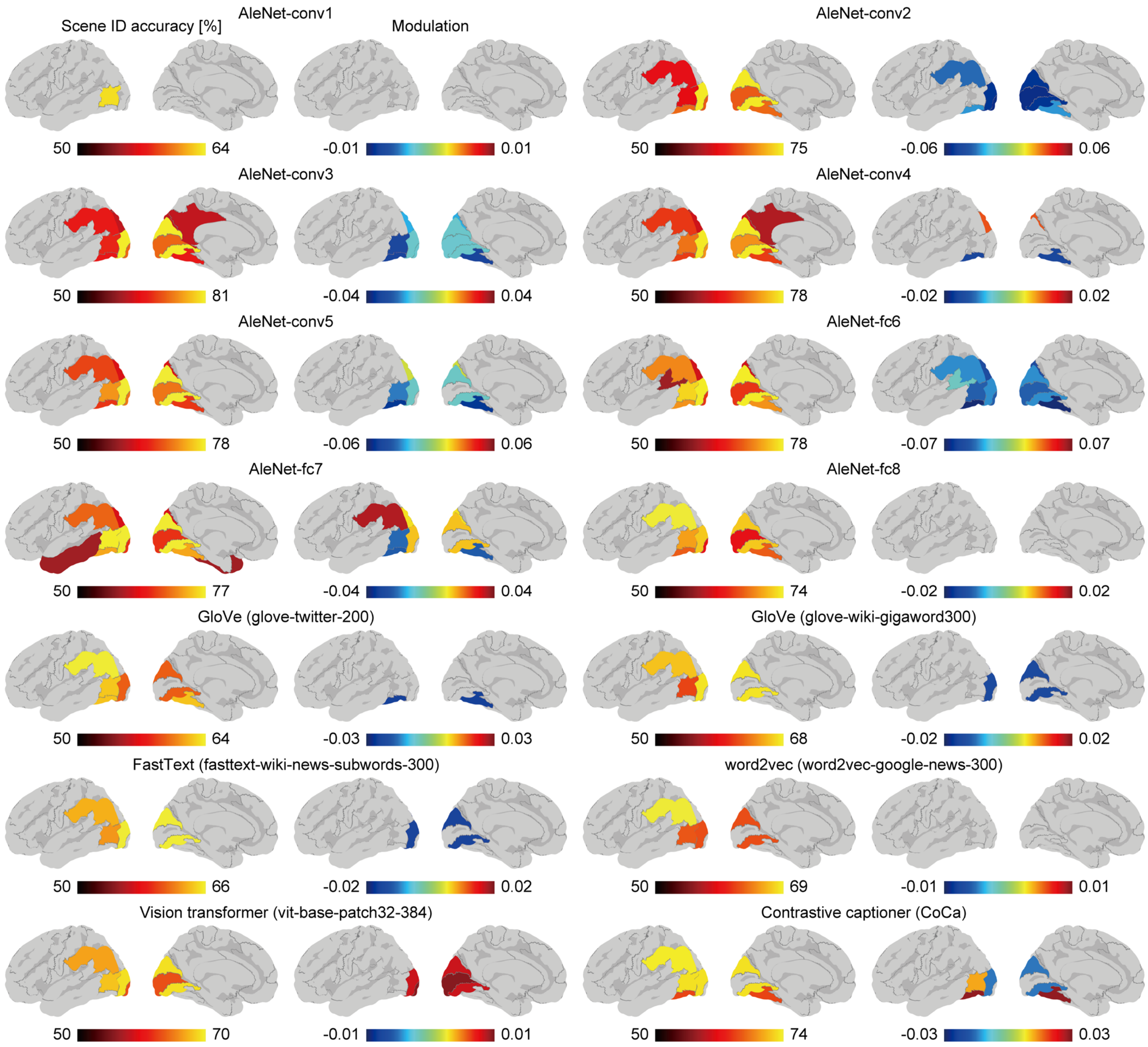


**Supplementary Fig. 8. Scene-identification accuracy and modulation in various latent spaces.**

In the same manner as in Fig. 4e and f, scene-identification accuracy and modulation of each cortical region were calculated from the virtual subject to be visualized with colour-coding on the normalized brain surface ($p < 0.05$, FDR adjusted among cortical regions for both scene-identification accuracies and modulations). Notably, the modulations were only calculated for the cortical regions that presented significant scene-identification accuracy. To encode images with a vision transformer (vit-base-patch32-384), latent vectors corresponding to the CLS token were extracted.

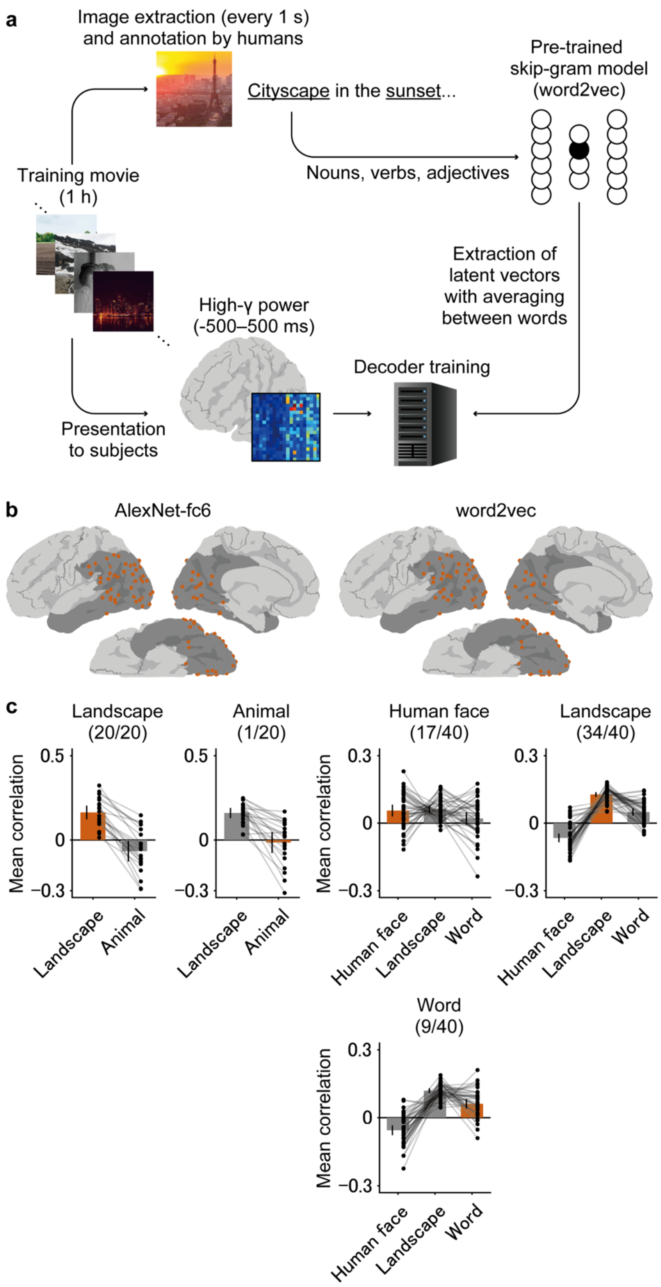


**Supplementary Fig. 9. Online task performed by EB01 using the AlexNet-fc6 and word2vec latent spaces.**

(a–c) EB01 participated in a session for a closed-loop online task using a word2vec latent space other than the session using the AlexNet-fc6 latent space shown in the main text. (a) A decoder used in the session was trained using the high-γ powers from ECoG signals segmented by 1-s time windows, while EB01 watched a 1-h movie. From the movie, 3,600 still images that were presented at the middle of the time windows were obtained to be annotated by humans. From the annotations, words (nouns, verbs, and adjectives) were extracted to be fed into a skip-gram model (word2vec) to acquire the latent vectors, which were later averaged within each image to create semantic vectors corresponding to the images. The decoder was trained exactly in the same way as that used in the session using the AlexNet-fc6 latent spaces. The interpolation factor ($\alpha$) was set to 0.5, which is the same value used in the session with the AlexNet-fc6 latent space. For details of the methods, see Methods and Fukuma et al., 2022[17]. (b) Electrode location of EB01 used for the session with AlexNet-fc6 and word2vec latent spaces is shown on the normalized brain, with colour-coding of red and blue denoting the electrodes on the left and right hemispheres, respectively. The cortical area marked with a darker colour denotes the regions where the subdural electrodes were located. (c) The average of 32 Pearson's correlation coefficients between the interpolated vectors (used to determine the feedback images) in a trial and the latent vectors for the target/nontarget instructions is shown as bars, with error bars denoting corresponding 95% CIs. For the session with the word2vec latent space, the instructions were "human face", "landscape", and "word". EB01 succeeded in controlling the feedback image with the word2vec latent space ($p < 0.001$, one-sided binominal test, $n = 120$) but failed with the AlexNet-fc6 latent space ($p = 0.437$, $n = 40$). Here, considering the results from EA02, who succeeded in controlling the CLIP latent space but also failed with the AlexNet-fc6 latent space, it was suggested that the difference in latent space affects the online performance. In addition, our previous study demonstrated that modulation in the word2vec latent space becomes significantly positive[17]. Hence, it was also suggested that positive modulation was one of the key factors in successful control of the feedback images.

**Supplementary Table 1. Demographics of the subjects**.

| Pt ID | Age | Sex | Performed task | | | Number of electrodes† | |
|---|---|---|---|---|---|---|---|
| | | | Image perception task* | Modulation task | Online task | Subdural electrodes | Depth electrodes |
| EA01 | 10–19 | M | ✓ ( 2 / 1 ) | | ✓ | 46 (23) | 12 (0) |
| EA02 | 30–39 | M | ✓ ( 3 / 1 ) | ✓ | ✓ | 60 (54) | 8 (0) |
| EA03 | 40–49 | M | ✓ ( 1 / 1 ) | ✓ | | 46 | 30 |
| EA04 | 20–29 | M | ✓ ( 2 / 1 ) | ✓ | | 36 | 12 |
| EB01 | 30–39 | M | ✓ ( 2 / 1 ) | ✓ | ✓ | 64 (62) | 0 (0) |
| EB02 | 10–19 | F | ✓ ( 3 / 3 ) | ✓ | ✓ | 88 (88) | 0 (0) |
| EB03 | 20–29 | F | ✓ ( 2 / 2 ) | ✓ | | 66 | 0 |
| EB04 | 10–19 | F | ✓ ( 1 / 1 ) | ✓ | | 76 | 0 |
| EB05 | 10–19 | F | ✓ ( 3 / 1 ) | ✓ | | 78 | 0 |
| EC01 | 20–29 | M | ✓ ( 2 / 1 ) | | ✓ | 100 (66) | 0 (0) |
| EC02 | 40–49 | F | ✓ ( 1 / 1 ) | ✓ | | 54 | 6 |

*The first and second values in brackets denote the number of training and evaluation sessions for the image perception task, respectively.

†Values in parentheses denote the number of electrodes used in the online task.
